# Diversity and Community Structure of Collembola (Hexapoda) across sites and microhabitats in the Bula Protected Area, a UNESCO-Listed Hyrcanian Forest at the Crossroads of Two Biodiversity Hotspots (Iran)

**DOI:** 10.1101/2025.05.20.650270

**Authors:** Thomas Tully, Masoumeh Shayanmehr, Elham Yoosefi Lafooraki, Mehdi Ghajar Sepanlou, Cyrille D’Haese

**Affiliations:** Sorbonne Université, UPEC, CNRS, IRD, INRA, Institute of Ecology and Environmental Sciences - Paris, IEES, Paris, France; Department of Plant Protection, Faculty of Crop Sciences, Sari University of Agricultural Sciences and Natural Resources, Sari, Iran; Centre for International Scientific Studies and Collaboration (CISSC), Ministry of Science, Research and Technology, Tehran, Iran; Department of Soil Science, Faculty of Crop Sciences, Sari University of Agricultural Sciences and Natural Resources, Sari, Iran; Département Adaptations du Vivant (AVIV), Muséum National d’Histoire Naturelle (MNHN) 45, rue Buffon 75005, Paris, France

**Keywords:** Alpha, beta and gamma biodiversity, Dead wood, Ecological niche, Forest Ecosystem, Indicator species, Mesofauna, Microarthropod community, Spatial heterogeneity, Species co-occurrence, Soil fauna, Springtail, Inter-specific occupancy-abundance relationship, biodiversity, evenness, species’ co-occurrence, springtail, interspecific occupancy-abundance relationship, dead wood, core, satellite, urban and rural species

## Abstract

**Background:** The Hyrcanian Forests of Iran, a unique 850 km stretch of deciduous forest along the southern coast of the Caspian Sea, are renowned for their exceptional biodiversity, high levels of endemism and ecological integrity. The northern part of this forest massif is part of the Caucasian Biodiversity Hotspot, while its southern fringe extends into the Iranian-Anatolian Hotspot. The recognition of these forests as a UNESCO World Heritage Site has focused global attention on their protection, primarily from human disturbance. While the diversity of vascular plants is well documented, other groups, such as soil-dwelling arthropods, remain under-explored. This study aims to fill this knowledge gap by conducting inventories in different sections of the Bula Forest, located in the eastern part of the Hyrcanian region, with a focus on Collembola, a potentially diverse but understudied group of soil fauna. The study was conducted to answer the following questions: Is the naturalness and degree of conservation of these forests reflected in the biodiversity of springtail communities? To what extent does the composition and diversity of these communities vary between different sites and microhabitats within these forests?

**Methods:** To answer these questions, 89 samples were collected over a two-year period from three different forest sites that had undergone contrasting forest management. Specifically, the study focused on three forest ecosystems, a preserved anthropized (planted) forest (PAF), a preserved natural forest (PNF) and a non-preserved natural forest (NPF), all located in the Bula protected area. Three types of microhabitats were sampled at each site, including moss, litter and soil, and dead wood. The springtails collected were identified and counted (approximately 3,000 individuals in total), allowing various biodiversity indices to be calculated for the communities as a whole, as well as by site and microhabitat. Soil samples were also analysed to compare soil properties between the three sites.

**Results:** The results reveal an impressive diversity of springtails, with 73 morphospecies identified, belonging to 39 genera and 12 families. Of these, 49 could be identified to species level. The estimated total species richness (gamma) ranged from 80 to 110 species. The analyses also revealed differences in species richness between sites, with two preserved sites (PAF and PNF) having a greater number of species compared to the non-preserved site (NPF), and in microhabitats, with greater diversity observed in dead wood, followed by soil and then mosses. A more detailed analysis of the communities showed a high degree of interspecific diversity in terms of abundance and distribution, without revealing strong patterns of ecological specialization for the majority of species. An analysis of co-occurrences showed that, in most cases species interactions appeared to be neutral. However, some pairs of species showed positive associations, while negative interactions were more marginal.

**Discussion:** The Bula Forest host particularly diverse Collembola communities. The results underline the importance of conserving these environments, which harbours a high diversity of springtails. Additionally, the results highlight the crucial role of dead wood as a substrate in supporting this biodiversity. The results also call for a more in-depth investigation of the factors influencing the presence, absence and abundance of species, as well as an exploration of the underlying causes of positive or negative species covariation.

## Introduction

The Hyrcanian forests in Iran are a unique and ancient hardwood ecosystem stretching along the southern coast of the Caspian Sea, extending from Azerbaijan to Iran’s eastern and northern provinces, including Guilan, Mazandaran, and Golestan (Akhani et al., 2010; Tohidifar et al., 2016). Dating back 25 to 50 million years, these forests served as major glacial refugia during the Quaternary glaciations, preserving numerous endemic species (Vakili et al., 2021; Sagheb Talebi et al., 2014; Tohidifar et al., 2016; Noroozi et al., 2019).

Rich in biodiversity, these mountain woodlands provide essential ecological and economic services, such as groundwater conservation, wildlife habitat, and erosion control (Haghdoost et al., 2011). Key tree species include *Fagus orientalis*, *Carpinus betulus*, and *Quercus castaneifolia* (Haghdoost et al., 2011; Mehrafrooz Mayvan et al., 2015a).

Despite their significance, the Hyrcanian forests have faced severe threats, losing about one third of their area since 1950, primarily due to human activities (Saeei, 1950; Tohidifar et al., 2016; Noroozi et al., 2023). Recognition as a UNESCO World Heritage site has raised global awareness of these forests, which are located in 15 regions, including Bula Forest (Tohidifar et al., 2016; Homami Totmaj et al., 2021; Vakili et al., 2021; Nourzad Moghaddam et al., 2018; Homami Totmaj et al., 2021).

Collembola (springtails), one of the earliest soil colonizers in the forests (Chahartaghi et al., 2005; Kuznetsova and Ivanova, 2020), play a crucial role in soil formation, nutrient cycling, and plant growth (Hopkin, 1997; Gange, 2000; Petersen, 2002; Filser, 2002). Local species richness and density of Collembola can vary a lot among sites and ecosystems (Paul et al., 2011; Bhagawati et al., 2021; Mehrafrooz Mayvan et al., 2022). Although specific diversity can reach a maximum value of around 100 species, it most often averages between 20 and 30 species (Potapov et al., 2023). Springtail density in soils vary widely, ranging from several tens of thousands to over a hundred thousand individuals per square metre; in high-latitude regions, these numbers can even sometimes exceed several hundred thousand per square metre (Potapov et al., 2023; Rusek, 1998; Hopkin, 2006; Harta et al., 2021).

There are about 9000 published Collembola species worldwide (Bellinger et al., 2024) while Iran’s share of this number was only 232 in 2020 (Shayanmehr et al., 2020) and reached 345 in 2025 (D’Haese et al., 2025; Shayanmehr et al., 2025). More than half of these species (250 species) were reported from Hyrcanian forests (D’Haese et al., 2025; Shayanmehr and Yahyapour, 2019). The number of published endemic species from these forests is on the rise (Shayanmehr et al., 2013; Yahyapour et al., 2020; Yahyapour et al., 2021; Yoosefi Lafooraki et al., 2020; Shayanmehr et al., 2013; Yahyapour et al., 2020; Yahyapour et al., 2021; Yoosefi Lafooraki et al., 2020; D’Haese et al., 2025; Shayanmehr et al., 2024). However, many species remain unknown in various areas of these forests, especially in the UNESCO-listed areas (Shayanmehr et al., 2020; Shayanmehr and Yahyapour, 2019; D’Haese et al., 2025).

Most studies on Collembola in Iran are qualitative faunistic surveys. Virtually no information is available on the diversity, density or structure of communities of these soil-dwelling arthropods. The only notable study is that conducted by Mehrafrooz Mayvan et al. (Mehrafrooz Mayvan et al., 2015a), which found that the depth distribution of Collembola resembled that of temperate forests. Seasonal dynamics differed, with the highest densities occurring in winter. Reduced moisture in summer is probably a major determining factor.

More broadly, the density and composition of a springtail community arise from complex interactions between movement, dispersal, biotic interaction, stochastic factors, and environmental filters such as drought and humidity, temperature, altitude, soil quality and chemistry (Petersen, 2011; Xie et al., 2022; Xie et al., 2025; Loranger et al., 2001; Holmstrup et al., 2018). Environmental filters – typically abiotic factors – predominantly shape the composition of communities at medium to large spatial scales (Ponge and Salmon, 2013; Potapov et al., 2020a). Biotic factors, such as the presence of predators or the intensity of intraspecific or interspecific competition, in contrast, generate significant spatial heterogeneity in springtail community composition at smaller spatial scales (Widenfalk et al., 2016). Dispersal may play a key role in determining the species pool at a large spatial scale, while species differences in individual mobility can influence local community composition. Ecological differences between species, such as differences in ecological niches, further modulate these species assemblages. For instance, generalist and highly mobile species occupy a wide variety of microhabitats, thereby reducing beta diversity, whereas specialist species reinforce differentiation between communities linked to distinct microhabitats.

Abiotic environmental filters function as niche selection mechanisms, alongside spatial and stochastic processes (dispersion, drift, and self-organization), with their relative importance varying markedly depending on the study scale. These effects interact in intricate ways, often forming a network of direct and indirect effects that complicates the derivation of clear, general predictions (Petersen, 2011).

In addition, various stochastic factors such as temporal and seasonal fluctuations in population size – which are difficult to quantify in the field – can amplify the spatial variability observed in community composition. Together, these sources of variation interact in complex ways, making ecological patterns inherently difficult to predict and interpret. This complexity underscores the value of experimental approaches that allow precise estimation of the impact of individual variables on the structure of springtail communities under realistic field conditions (Holmstrup et al., 2018; Lindberg et al., 2002).

Without claiming to untangle all or part of this complex network of causal relationships, a series of samplings of springtail communities was conducted as part of this study. The objective of this sampling was to better understand the diversity, composition and spatial structure of Collembola communities in the Hyrcanian Forests. More specifically, sampling took place in the Bula Forest, a pilot site within the UNESCO-listed ancient Hyrcanian Forests, located in Mazandaran Province in the eastern part of Iran’s Hyrcanian region. We surveyed three distinct sites with different management history – natural preserved, anthropized preserved, and non-preserved natural forest – sampling three microhabitats in each (litter, dead wood, mosses). This experimental protocol provides a sampling design for measuring spring springtail biodiversity at different spatial scales, among sites that are few kilometres apart and among three microhabitats.

This sampling enabled us to identify 73 morpho-species, belonging to 39 genera and 12 families. Of these, we were able to accurately identify 49 species. The list of these species and their precise description, illustrated with photographs, is the subject of a first publication, in which we have also summarized the specific diversity known to date of Hyrcanian forests (D’Haese et al., 2025).

In this study, we employed these previously compiled species inventories to assess biodiversity beyond a basic overall measure of species numbers. More precisely, we estimated several levels of biodiversity: gamma biodiversity (γ), corresponding to the pool of species observed in the study area; alpha diversity (α), measuring the mean specific diversity of each sampling site; and several dimensions of beta diversity (β), indicating heterogeneity between sites, and between microhabitats.

We examined patterns of diversity at several nested spatial scales: (1) at a broad scale, across the entire study area; (2) at the intermediate scale of three distinct sites, each with a different forest management history and located within a few kilometres of each other; and (3) at a finer scale, across three contrasting microhabitats present at the three sites. This multiscale approach allows us to assess how local microhabitat variation – even within a single site – can support distinct Collembola communities (Berg, 2012).

Our dataset also allowed us to investigate whether springtail communities comply locally with the general ecological interspecific occupancy-abundance relationship (AOR): the abundance of a species is generally positively correlated with its presence or occupancy. In other words, species with the widest distribution, i.e. those present in a large number of habitats, are also often the most abundant locally (Gaston, 1996; Brown, 1984; Blackburn et al., 2006; Gaston et al., 1997). This positive relationship is expected especially when the communities are composed of “core” species or “common” species that are both abundant and widespread and of “satellite” species, i.e. “rare” species that occupy a small proportion of sites and have also a small local population sizes (Hanski, 1991; Avolio et al., 2019). However, communities can also include two other types of “rare” species. First, “rural” or “spared” species that are widespread but occur with consistently small local population sizes. Second, “urban” or “restricted” species that reach high local abundances but are confined to only a small subset of the available habitats (Söderström, 1989; Avolio et al., 2019; Collins et al., 1993; Hanski, 1991).

The species-specific composition of a local community is not only determined by dispersal and environment filtering but can also be determined the composition of the community through biotic interactions (competitive exclusion for instance, Hishi et al., 2022; Caruso et al., 2013). Interspecific interaction between Collembola that share a similar niche can be complex, and sometime asymmetrical (Thakur et al., 2017; Theenhaus et al., 1999). We finally searched for indicators of positive or negative interactions between species, whether direct or indirect, based on their co-occurrence patterns in the field.

To our knowledge, these last two issues have been studied only to a very limited extent, if at all, in springtails in the wild.

In summary, our study aims to address the following research questions:

1. Do the Hyrcanian forests – internationally recognized for their ecological value and located at the intersection of two biodiversity hotspots – harbour particularly diverse Collembola communities?
2. How does Collembola diversity vary among different forest sites and across distinct microhabitats (e.g. soil leaf litter, dead wood, moss)? Specifically, we measured and compared, diversity (alpha, beta and gamma) at the community level, as well as within each forest sites and microhabitat types. This analysis allows us to determine whether springtail communities exhibited strong microhabitat specialization (indicating dominance by specialized species or indicator species) or whether their spatial structuring is more diffuse, suggesting the prevalence of generalist species across broader spatial scales.
3. Are Collembola community evenness homogeneous across sites and microhabitats? By examining community evenness, we moved beyond simple species counts to quantify how relative abundances are distributed among species. This provides a more nuanced understanding of community structure and potential differences between sites or microhabitats. While our experimental design limits our ability to explicitly link forest management to community composition, we hypothesize that natural and protected forests may harbour greater species diversity or more even communities compared to more anthropized forests, reflecting the role of conservation measures in maintaining balanced and diverse communities.
4. Which species dominate in terms of local or global abundance and occupancy? How do abundance and occupancy vary and co-vary among species? Do these variables covary positively, consistent with the general abundance-occupancy relationship observed in some ecological communities or are the collembola communities also composed of “urban” and “rural” species?
5. What are primary drivers shaping the local Collembola community compositions? We investigate the relative contributions of dispersal limitations (expected to drive differences among sites), environmental filtering (expected to drive differences among microhabitats), biotic interactions, particularly interspecific interactions among Collembola species (e.g. competition or facilitation). Communities dominated by competitive interactions should exhibit negative co-occurrence patterns, reflecting species segregation. Conversely, positive co-occurrence may indicate shared microecological preferences or mutualistic interactions.

## Material and methods

### Sampling Protocol

The samplings were performed in the Bula Forest, which has a semi-humid climate and is located in the south of Do-Dangeh section of Sari (Mazandaran Province, Iran) and in the north of Khere-Nero (part of Alborz) mountain range (Figure 1). The Bula Forest has significant elevation differences in its various parts. The 3907 hectares of this protected area are covered with dense forests that are under the management of the Environmental Protection Organization of Iran (EPOI). The main vegetation types covering these territories are lowland deciduous forests, submontane and montane deciduous forests, Juniperus and Cupressus woodlands, cultural landscapes and artificial forests, riverine and valley forests (Akhani et al., 2010).

**Figure 1.**
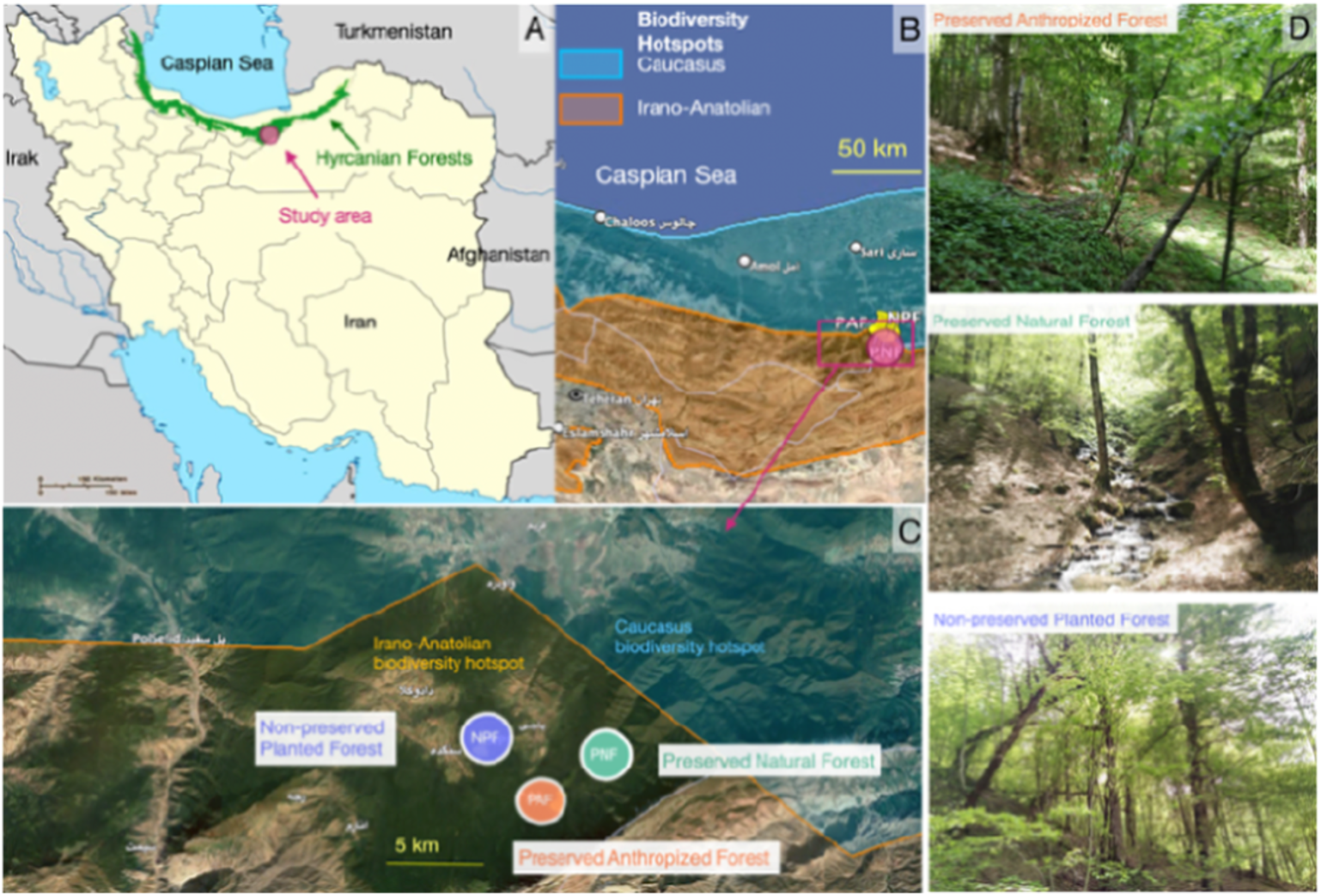
The general location of the study area located in the Bula Forest in northern Iran. Mazandaran is marked with a red circle on the Hyrcanian Forest (green) on a general map of Iran (A). A closer look (satellite maps B & C) shows the exact location of the three sites, a preserved anthropized forest (PAF), a preserved natural forest (PNF) and a non-preserved natural forest (NPF). These sites are formally located in the Irano-Anatolian Biodiversity Hotspot, but only a few kilometres from the border of the Caucasian Biodiversity Hotspot (Hoffman et al., 2016). The images on panel D show the general appearance of the three sites.

To broaden the chances of observing different species, and to provide a first assessment of the spatial variability of the Collembola communities, we took samples from three sites at various altitudes and undergoing contrasted forest management (Figure 1). The first site, at a higher elevation (1670 m a.s.l.), is pristine and undisturbed, representing a preserved natural forest (later called ‘PNF’). The second site, at a slightly lower elevation (1600 m a.s.l.), has been reforested as part of conservation efforts, classified as a preserved anthropized forest (“PAF”). Both of these sites are managed by the Environmental Protection Organization of Iran (EPOI), fenced off, and restricted from public access. The third site, situated at a lower elevation (1425 m a.s.l.), is open to the public and classified as a non-preserved forest (‘NPF’). The three selected sites therefore differ in terms of forest management history, but also in terms of altitude and probably of many other covariates that were not included in our analysis. They were chosen in order to broaden the sampling to include different ecological conditions, but the sampling design does not allow for a systematic study of the potential effect of altitude or forest management on springtail communities (lack of replication).

Within each site, three types of microhabitats (dead wood, moss, soil and leaf litter) have been collected from various points, at about 30 steps apart (∼20m). This was done twice, in late April 2021 and in early June 2022. As a total, within each site, three samples of soil and leaf litter, three samples of moss, and four samples of dead wood were collected in 2021. In 2022, with the protocol now more established, we were able to take more samples: five per site for moss and dead wood, and ten per site for forest soil. This amounts to a total of 90 samples (30 samples per site), 89 of which could be analysed (no Collembola was found in one forest soil sample in 2022). It should be noted that sampling was conducted in spring, which is generally considered a favourable period, although not necessarily the most favourable one (Mehrafrooz Mayvan et al., 2015a; Mehrafrooz Mayvan et al., 2022). Consequently, part of the community diversity may have been underestimated due to seasonal fluctuations. As no sampling was carried out in autumn or winter, it is possible that several species that are particularly abundant or detectable during these seasons were not recorded.

The soil and leaf litter samples were taken using a 5 cm-diameter soil corer to a depth of 10 cm, allowing us to sample both the organic surface horizons (O_Litter_, O_Fragmentation_ and O_Humus_; approximately 0-3 cm) and the underlying organo-mineral A_H_ horizon (∼3-10 cm), following Mehrafrooz Mayvan et al. (2015a). Moss and dead wood samples were collected in 500 mL plastic containers. The samples were kept separate from each other, tagged and transported in the afternoon to the laboratory and the soil fauna was then extracted using Berlese-Tullgren extractors (D’Haese et al., 2025; Valle et al., 2023). The samples were kept in the funnels for one to two weeks until all Collembola had been fully extracted. During this period, the samples were checked every three to four days, and any Collembola present were removed. A 60 W incandescent lamp was placed above each soil sample, although the temperature was not recorded.

In order to characterize the soil physicochemical properties at the sampled sites and to investigate possible differences between the three sites, we collected three soil samples per site at a depth of 0-10 cm, as for the soil samples for Collembola, resulting in a total of nine samples. All samples were collected on the same day. The soil analysis was conducted at the Soil Laboratory of Sari University of Agricultural Sciences and Natural Resources (SARU) to assess both physical and chemical properties (Figure 2). Physical properties included soil texture, determined by the percentage of clay, silt, and sand by hydrometer method (Gee and Bauder, 1986). Chemical properties included % organic carbon, determined by titration and coloric methods (Nelson and Sommers, 1996), and pH, measured by electrochemical methods using pH meter (Rhoades, 1982).

**Figure 2.**
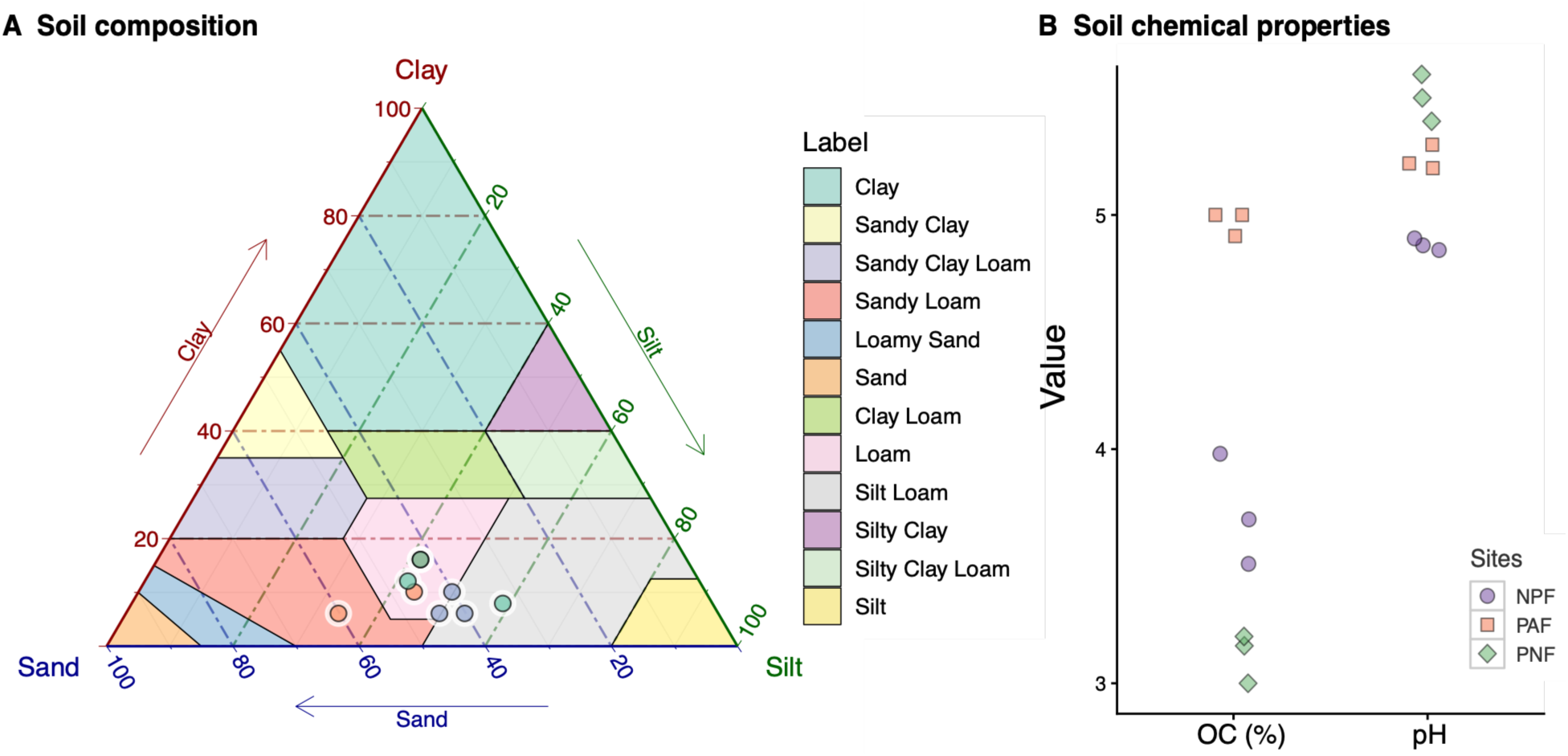
Soil composition was investigated by estimating the percentages of clay, sand and silt for three replicate samples within each of the three sites (panel A). We also report on panel B the measurements of two soil chemical properties - % of organic carbon (OC) and pH - for the nine soil samples analysed. NPF for Non-preserved Natural Forest, PAF for Preserved Anthropized Forest and PNF for Preserved Natural Forest.

### Collembola Sorting and Species Identification

For each sample, the specimens were examined under a dissecting microscope, sorted by morphospecies and, for each morphospecies, the number of individuals for each morphospecies was determined. The morphospecies were sorted according to shape and colour pattern. For species identification, some specimens were cleared in 10% KOH (for light specimens) and Nesbitt’s solution (for dark specimens) and mounted in Hoyer’s medium (Mari-Mutt, 1979). They were examined under a Nikon’s ECLIPSE E600 microscope. Species were identified on the basis of morphological taxonomic characters described by Fjellberg (Fjellberg, 1998; Fjellberg, 2007), Potapov (Potapov, 2001), Jordana (Jordana, 2012) and Bellinger et al. (Bellinger et al., 2024). All materials are kept at the National Natural History Museum in Paris, France. The list and description of the species identified are published in two accompanying articles (D’Haese et al., 2025; Shayanmehr et al., 2024) and will not be discussed in detail here.

When the exact species name could not be determined, the morphospecies were named with their genus names followed by ‘sp.’. The species names ‘sp.01’, “sp.02’… were used to designate morphologically different morphospecies belonging to the same genus (D’Haese et al., 2025). Some Neelipleones were present in the samples, but given their low numbers and rarity in some of the samples, compared to their expected density in such microhabitats, it has been assumed that most of them escaped the attention of the examiners during the sorting of the samples, especially from the first session (2021), and therefore we did not include them in some of the quantitative analyses (PCA). Similarly, some Symphipleones could not be accurately identified and are therefore not included in these quantitative analyses either. On the other hand, a great deal of identification work was carried out on the Neanuridae, where almost all the specimens could be identified to species (D’Haese et al., 2025).

### Statistical Analysis

#### Number of Specimens per Sample and Alpha, Beta and Gamma Species Diversity

All the statistical analyses were performed using R (The R Core Team, 2025) and we used the *ggplot2* package to construct the figures (Wickham, 2009).

Before carrying out the analysis, we deleted the data where the springtails could not be assigned to an exact species or morphospecies. This includes certain damaged or juvenile specimens.

We carried out our analysis using a table with 479 rows, from which we created a community data frame with 73 columns (different species) and 89 entries (we had 90 different samples, but in one of the samples no Collembola could be identified), using the function *matrify* from the R package *labdsv* (Roberts, 2023).

Biodiversity analyses were performed on three spatial scales. On a global scale, all samples were used to estimate alpha, beta and gamma diversities; on a microhabitat scale, we compared these diversities between the three microhabitats, namely dead wood, forest litter and moss; at the forest sites scale, we compared these diversities between the three sampling sites. For these analyses, the presence or absence of species was taken into account (number of species), but not their abundance, which was taken into account in subsequent analyses.

The mean number of Collembola per sample and the mean number of species per sample (local species richness or alpha (α) diversity) were analysed using quasi-Poisson generalized linear models (*glm* function), starting first with models that included the explanatory variables ‘site’ and ‘microhabitat’ and their interaction. These models were then simplified by removing the non-significant interactions (Faraway, 2016) and the simplified models were used to estimate the mean numbers of specimens or species per sample along with their 95% confidence intervals (Figure 3). Alpha diversity was estimated for the whole community and also for each forest site and each type of microhabitat.

**Figure 3.**
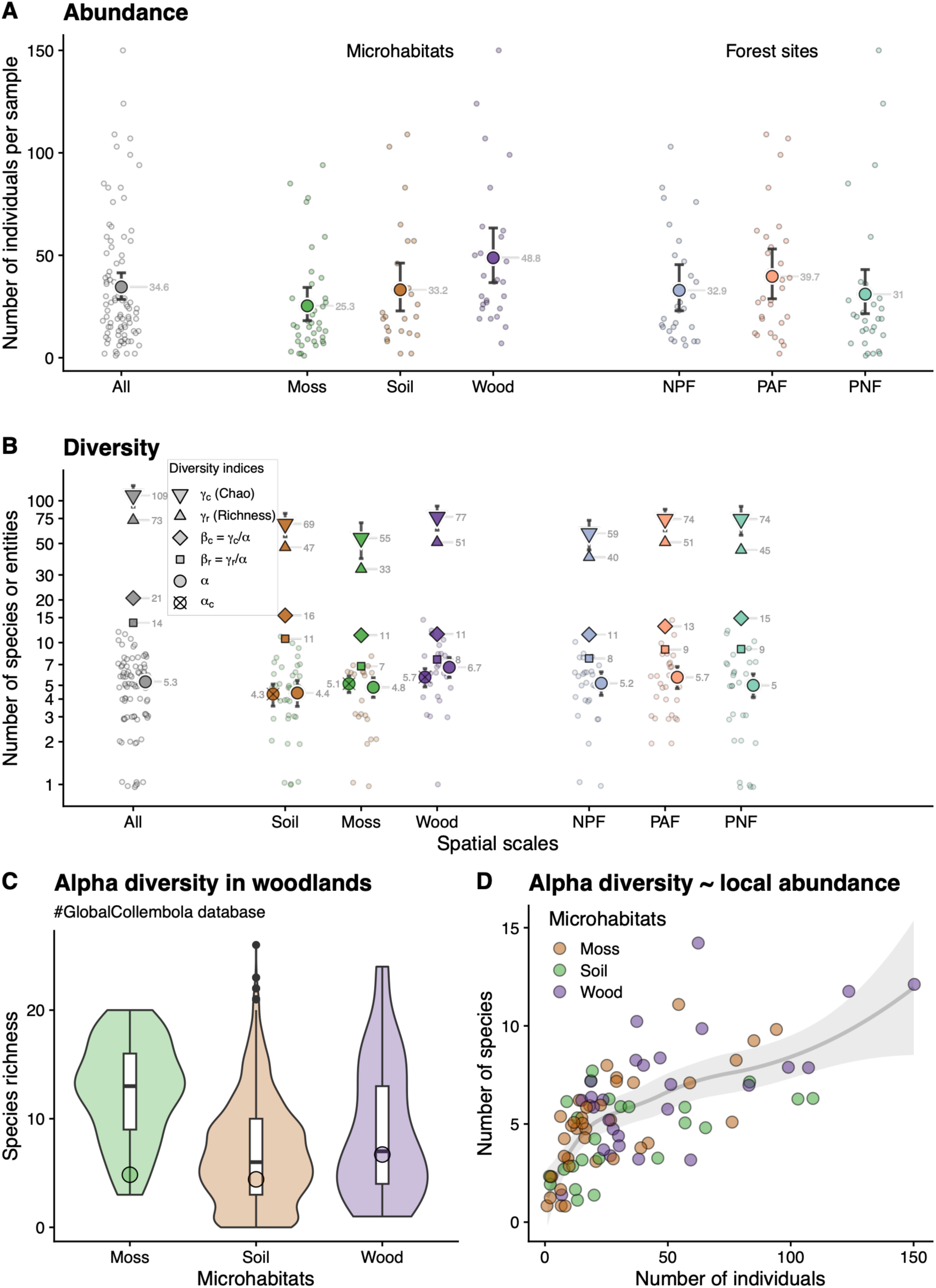
Number of Collembola per sample (A, abundance) and diversity (B) for all the samples combined together (All) or as a function of microhabitat types (middle) or of forest sites (right). Panel C displays the distribution of alpha diversity values from the #GlobalCollembola database, sourced from moss, forest litter (soil), and rotten wood in forest ecosystems. Coloured discs represent our average alpha diversity values, enabling a direct comparison with the global distributions. Panel D shows the relationship between species richness per sample and the corresponding number of individuals. The raw numbers of Collembola or species per sample are plotted on panels A & B (small dots), together with the mean and associated 95% confidence intervals derived from the simplified GLM quasi-Poisson models used to provide these estimates. On average, the number of individuals and species per sample is similar for soil and moss samples, but higher for wood samples. Panel B displays alpha diversity (α is the number of species per sample and α_c_ is the abundance-corrected diversity, estimated for an average abundance of 35 individuals per sample), along with two gamma diversity measures (top triangles: Chao-estimated total diversity, shown with ±SE error bars and observed total species count), and two beta diversity estimates (middle), derived from these gamma measures. The different β give the effective numbers of distinct communities. Note that the y-axis of panel B is plotted on a logarithmic scale.

Gamma diversity (γ), or species richness at a global scale, was measured in two different ways: firstly, by simply counting the number of different species observed on a global scale, and secondly, by estimating this diversity using the asymptotic values towards which the species accumulation curves converge. Species accumulation curves were plotted using the *vegan* (Oksanen et al., 2026) and *BiodiversityR* (Kindt and Coe, 2005) R packages. To compare species richness among sites and among microhabitats, we plotted species accumulation curves for the three sites and for the three microhabitats independently (Figure 4). Gamma diversity was measured as the expected species richness of the entire community (all samples pooled) or as a subset of the whole community (per microhabitats or per sites, as for α). We used the *diversityresult* function from the *BiodiversityR* package with various methods to compute different gamma diversity estimators (Chao, bootstrap and first and second order jackknife).

**Figure 4.**
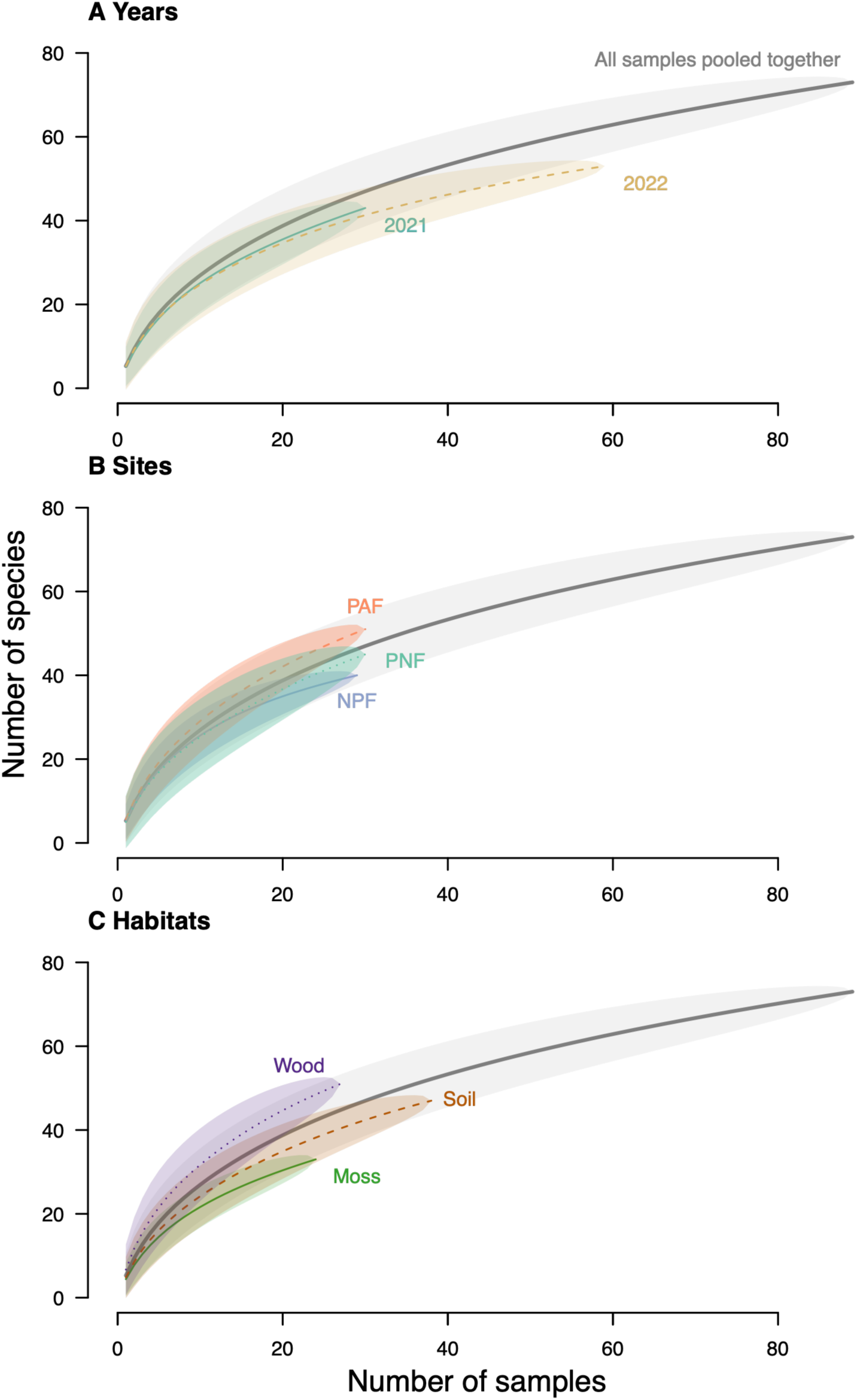
Species accumulation curves estimated using all different samples (grey lines) or different subsets of samples. Panel A shows the two years separately. Panel B allows comparison of the accumulation curves estimated for the three sites independently, namely Preserved Anthropized Forest (PAF), Preserved Natural Forest (PNF) and Non-preserved Natural Forest (NPF). Panel C shows the accumulation curves for the three microhabitat types (dead wood, soil or moss).

Measuring beta diversity (β) is less straightforward, as this term encompasses a wide range of concepts and phenomena that can have very different biological meanings (Koleff et al., 2003; Tuomisto, 2010b; Tuomisto, 2010a; Tuomisto, 2011). Here, following several authors’ recommendations (Tuomisto, 2011; Marcon, 2015), for the sake of simplicity and to enable direct comparisons with other diversity estimates, we have chosen to measure beta diversity as the ratio of gamma diversity to alpha diversity (multiplicative partitioning, β=γ/α). We used both the species richness and Chao’s estimator to get two estimates of beta diversity (β_r_ and β_c_ respectively). With this definition, beta can be understood as the ‘effective number of distinct communities’ which is the number of distinct communities required to reach the measured or estimated gamma diversity (Jost, 2007).

We compared our alpha biodiversity measurements with species richness distributions extracted from the #GlobalCollembola database (Potapov et al., 2024). Specifically, we focused on diversity data from three woodland microhabitats: moss (n = 83), litter (n = 2774), and rotten wood (n = 38), where specimens were collected using drying methods (Figure 3C).

#### Evenness

To account for evenness in biodiversity comparison between sites and microhabitats, we used Rényi’s diversity profiles (Rényi, 1961). These profiles allow the comparison of species richness and evenness of communities at a glance (Figure 5). At scale 0, the estimated diversity reflects the species richness on a logarithmic scale. At scale 1, it corresponds to the Shannon index. At scale 2, it shows *-ln(Simpson index)*. And for scale *inf*, *-ln(Berger-Parker index)* is plotted (Loreau et al., 2010). A flat profile indicates that the different species have the same evenness, while a declining profile is a sign of a community where species are unevenly distributed (Kindt and Coe, 2005). We used a stratified bootstrapping of the community matrix to estimate 95% confidence bands for the Rényi’s profiles.

**Figure 5.**
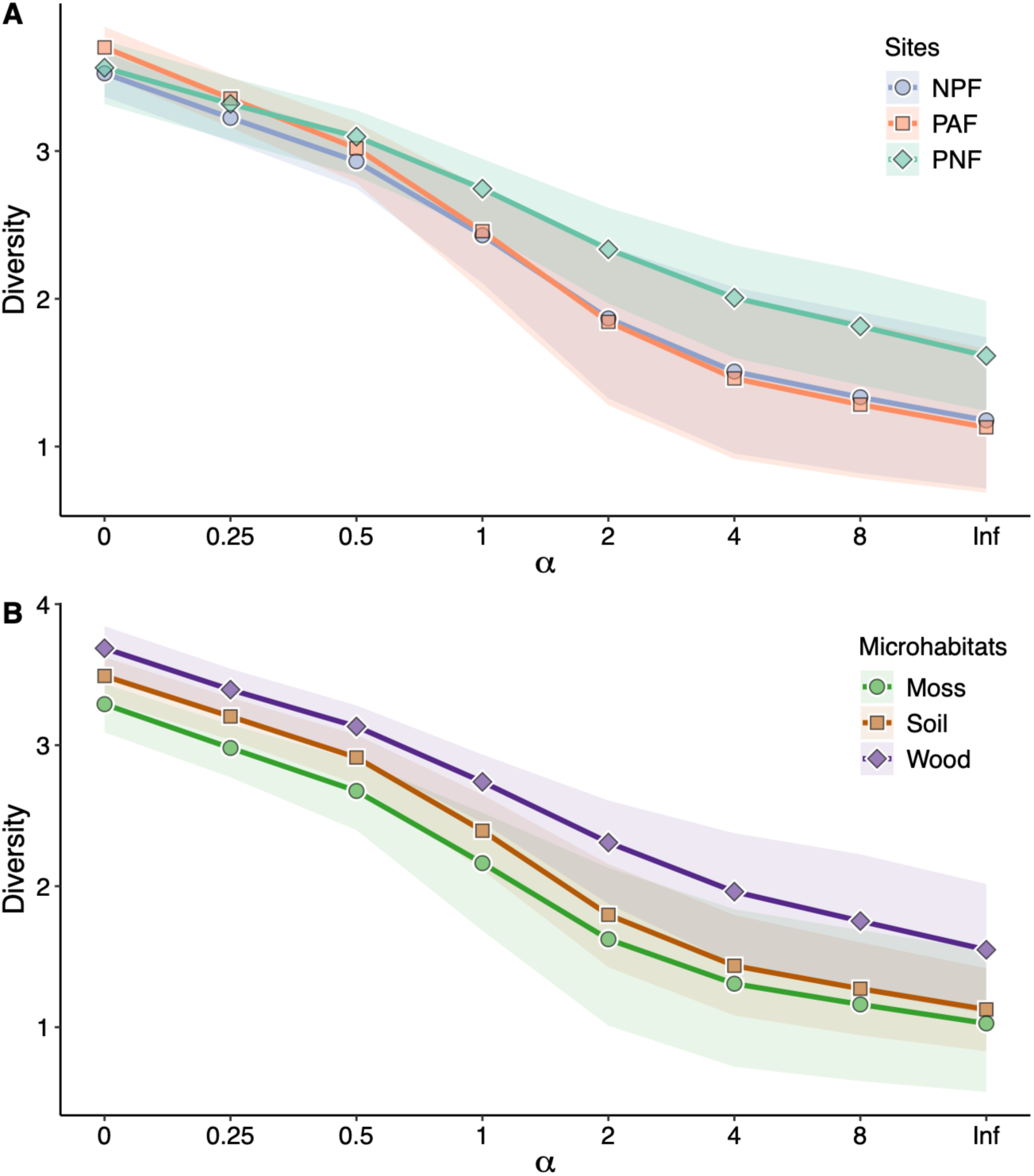
Rényi diversity profiles with 95% confidence bands. Comparison of the three sites (A) and of the three microhabitats (B). The scale parameter (α) changes the influence of species richness and evenness on the measurement of diversity. For a scale of zero, the relative abundances are not taken into account and the diversity is equal to the *ln* of species richness. For a scale of 1, diversity corresponds to the Shannon index. For a scale of 2, diversity equals *-ln(Simpson’s index)* and at α=inf diversity equals *-ln(Berger-Parker’s dominance index)*.

#### Species Abundance and Occupancy

To identify the species that were found to be the most abundant, or the most ubiquitous among our samples (high occupancy), we sorted the species according to their total abundance – measured as the total number of individuals summed across all the samples, Figure 3A – and indicated alongside their occupancy, which was determined as the proportion of samples in which each species was found (Figure 3B). These proportions, and their 95% confidence intervals, were determined, using a generalized linear model (GLM) for binomial variables.

The interspecific occupancy-abundance relationship was examined graphically at two different scales. At a global scale (Figure 7A), abundance was defined as the total number of individuals for each species across all samples, as shown in Figure 6A. At a local scale, abundance was calculated as the average number of individuals in samples where the species was present, focusing only on those specific samples (Figure 7B). We used these graphics to reveal the dominant Collembola species and to explore how the occupancy and abundance relate to each other in these communities.

**Figure 6.**
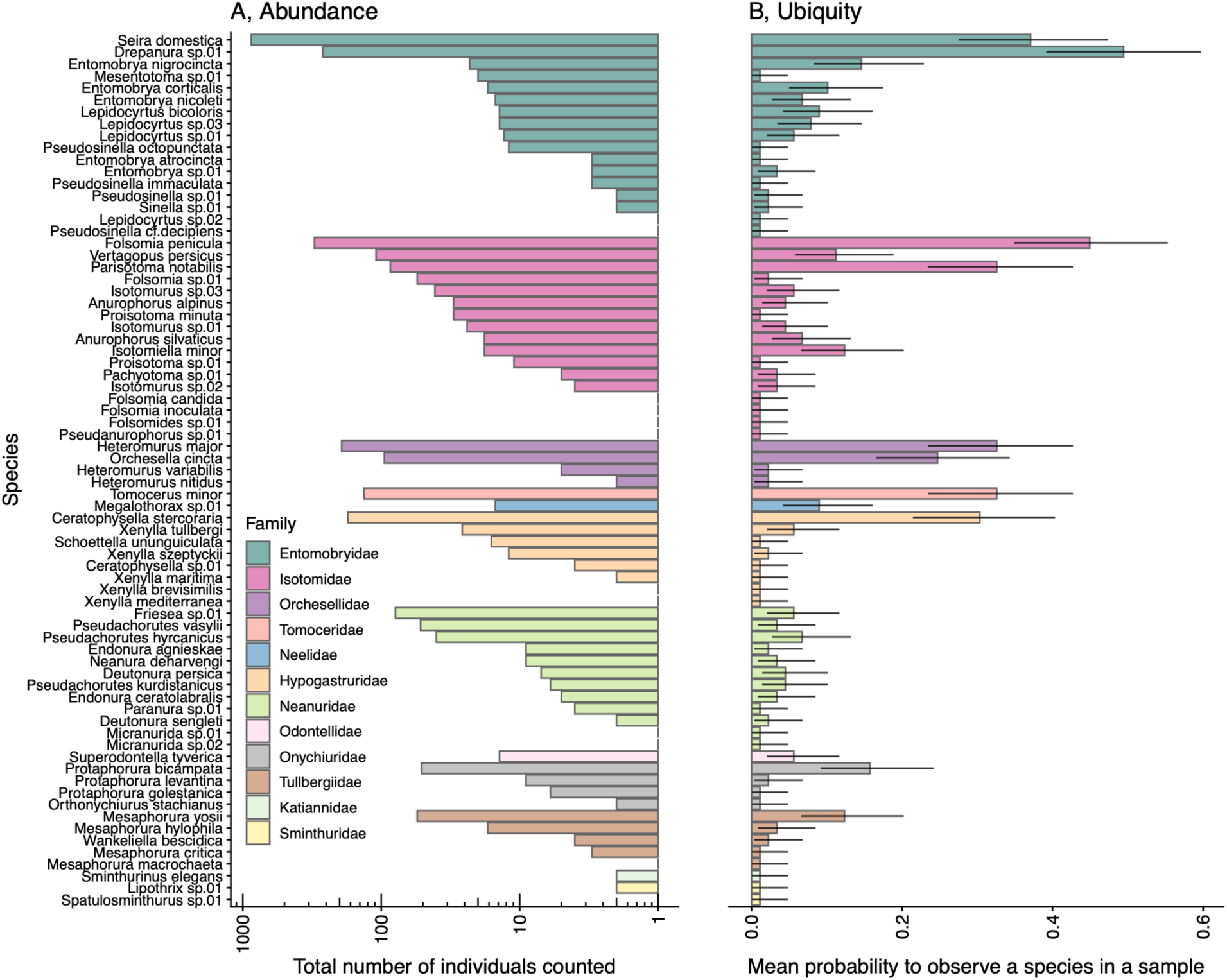
Panel A shows the total number of specimens collected for each morphospecies identified (on a logarithmic scale). It corresponds to a rank-abundance curve, but the species have been grouped by families, which are highlighted in different colours. Panel B shows, for each species, the proportion of samples in which we observed the species. The grey bars represent the 95% confidence intervals for each estimated proportion. The species are ordered from top to bottom by order, family and decreasing total abundance.

**Figure 7.**
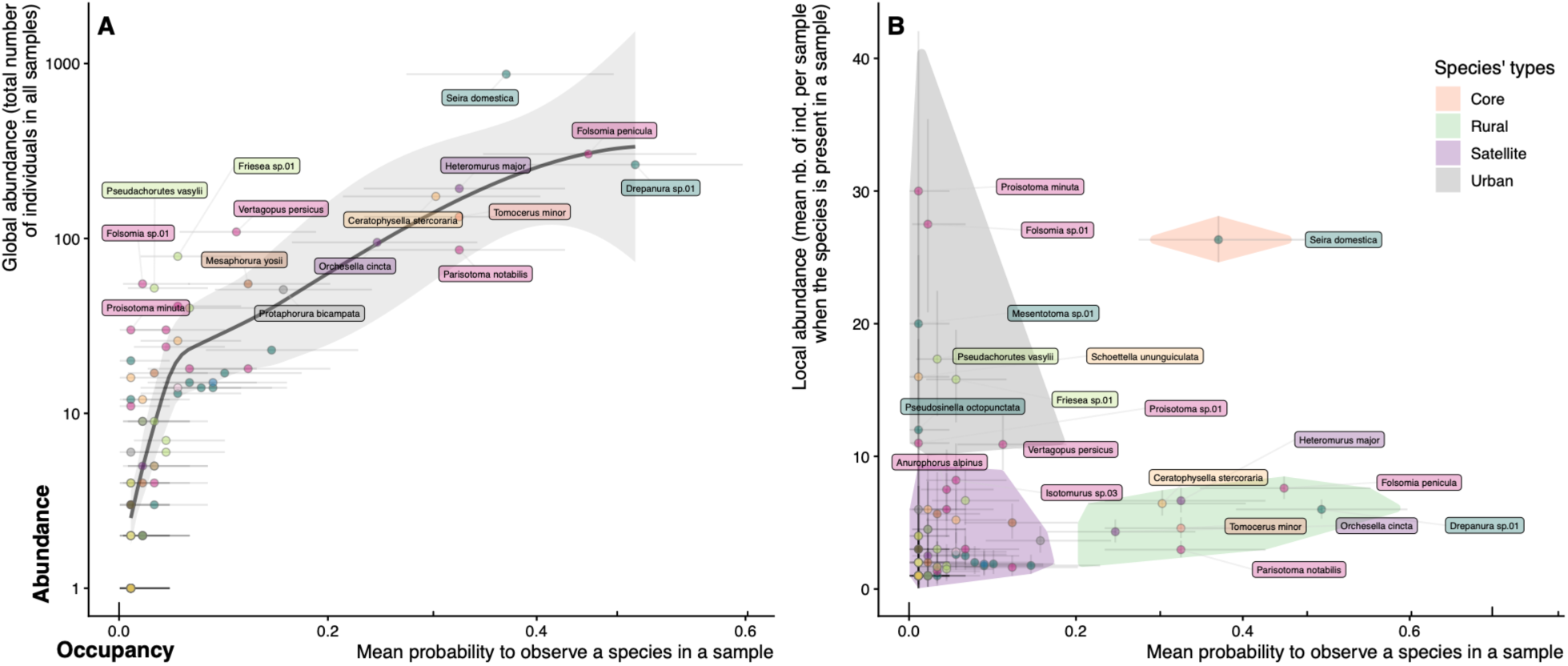
This figure shows the relationship between abundance and occupancy, expressed as the proportion of samples in which the species was observed. The abundance has been measured either by summing, per species, all the individuals collected in the different samples (global abundance, A, log10 scale), or by averaging the number of individuals in the samples in which the species is present (local abundance, B). The colours represent the different families (as for Figure 6) and the error bars represent the 95% confidence intervals estimated by the GLM models used to measure the occupancy (binomial model) or local abundance (Poisson model). We fitted a LOESS curve to the data on panel A to graphically reveal the relationship between the two variables and to show, in particular, the non-linearity between log(abundance) and occupancy without forcing a parametric model. On panel B, species have been grouped into four types depending of they mean local abundance (below or above 10 ind/sample) and mean occupancy (present in more or less than 20% of the samples, see main text in the introduction to a definition of the four types of species).

#### Collembola Communities, Ecological Specialization and Species Co-occurrence

We used a principal component analysis to study how the Collembola communities are organized, using the *FactoMineR* and *factoextra* libraries (Lê et al., 2008; Kassambara, 2017). We used this tool to look for any emerging structure in the samples according to the type of microhabitat or the sampling site. In this analysis, we excluded the species that were present in very few samples and retained only those that were present in at least 10% of the samples (13 species). We also performed a non-metric multidimensional scaling (NMDS) analysis based on Bray-Curtis dissimilarities using the *vegan* package (Oksanen et al., 2026) to compare the community composition among sites and microhabitats. We further used a analysis of similarity (ANOSIM) which uses a permutation procedure to look for statistically significant differences between sites or microhabitats.

We then adopted a graphical approach combined with an indicator species analyses, using the function *multipatt* from the R package *indicspecies* (De Cáceres and Legendre, 2009), to test for ecological preferences among the three microhabitats for the 25 most common species (Figure 9). We used this analysis to identify species significantly associated with one or two of the three microhabitats, and set the significance level to α = 0.005 to account for multiple testing across species.

Finally, we explored our dataset to study whether the presence of one species in a sample was linked to the presence or absence of another species (Figure 10). To do this, we used the *cooccur* R package (Griffith et al., 2016), which is designed to classify the pairs of species as representing random, positive or negative associations. For example, a pair of species is classified as positively associated if the observed frequency of co-occurrence is significantly greater than the expected co-occurrence.

## Results

### Soil quality

Despite differences in forest management practices, the three sites exhibited comparable soil composition (Figure 2A). The soil pH differed significantly among sites (p<0.01), although the overall range of variation was relatively narrow, with values spanning from 5.0 to 5.5. Similarly, the levels of soil organic carbon content differed between sites (p<0.01), with soils from the preserved anthropized forest showing the highest concentrations of organic carbon (Figure 2B).

### Species Identified

From the 89 soil, moss and wood samples, we sorted 3095 specimens of Collembola, which we then classified into 75 different morphospecies groups. We removed two of these morphospecies from the diversity analysis because they contained mainly juvenile individuals or individuals that were not distinct enough to be attributed to anything other than a genus (labelled ‘*sp*.’). Furthermore, we could not be sure that the individuals in these groups belonged to two or more species that were different from the others.

Of the remaining 73 morphospecies, 49 (67%) were identified at species level. The remaining 24 morphospecies were identified at genus level (labelled *sp.01*, *sp.02*, etc.) and were visually distinct enough from each other to be considered separate morphospecies. We found many *Tomocerus* juveniles in our samples. Given that we only identified *T. minor* in the adults, we classified all of these juveniles as *T. minor*.

The identified specimens belong to four orders and 12 families - Entomobryomorpha (Entomobryidae, Isotomidae, Orchesellidae, Tomoceridae); Symphypleona (Katiannidae, Sminthuridae); Poduromorpha (Hypogastruridae, Neanuridae, Odontellidae, Onychiuridae, Tullbergiidae); Neelipleona (Neelidae) - and 39 genera. The full list of species and their descriptions can be found in another article (D’Haese et al., 2025).

### Collembola Abundance and Alpha Diversity in the Samples

We found an average of 34.8 specimens (median 24, range 1–150, Figure 3A) and an overall mean alpha diversity of 5.3 species per sample (median 5, range 1–14, Figure 3B). Using quasi-Poisson generalized linear models, we found that the site*habitat interactions were not significant for either Collembola abundance (χ^2^_4_=8.7, P=0.069) or alpha species diversity (χ^2^_4_ = 1.5, P = 0.82).

Simplified models without interaction terms revealed that neither the total number of specimens nor the number of species per sample (α diversity) differed significantly among the three sites on average (abundance: χ^2^_2_ = 1.27, P = 0.53, alpha diversity: χ^2^_2_ = 1.21, P = 0.54). However, both metrics varied among microhabitat types (abundance: χ^2^_2_ = 8.87, P = 0.012, alpha diversity: χ^2^_2_ = 11.7, P = 0.003). Specifically, dead wood samples harboured on average more springtails and a greater number of species than either moss or soil samples, with no significant difference observed between the latter two microhabitats (Figure 3A & B). Our alpha diversity measures are fairly close to the median values from the #Globalcollembola database for soil and wood microhabitats, but lower than the diversity levels typically observed in moss-type microhabitats (Figure 3C).

Since alpha diversity increased with abundance (Figure 3D), we ran a third set of models to determine whether alpha diversity differed between sites and microhabitats when accounting for abundance in the model. As with the previous models, the interaction between site and microhabitat remained nonsignificant (χ^2^_4_ = 0.91, P = 0.98). The simplified model indicated that alpha diversity did not vary among sites (χ^2^_2_ = 0.79, P = 0.67), and the previously observed difference between microhabitats became only marginally significant (χ^2^_2_ = 5.8, P = 0.053). This suggests that part of the observed variation in diversity among microhabitats can be attributed to differences in Collembola abundance across these microhabitats (Figure 3B).

Conversion of the number of specimens per sample into density, that is to say in number per square metre, was only possible for soil samples for which we know the sampling surface (Ø 5 cm). This resulted in a mean density of 12450 springtails.m^-2^ (95% confidence interval: 8900–16820 ind.m^-2^).

### Species Accumulation Curves and Gamma and Beta of Diversity

The estimated extrapolated diversity (overall gamma diversity) is about 100 species for the three sites and three microhabitats combined together (Table 1 and Figure 4B), although we have clearly under sampled certain groups of Collembola (D’Haese et al., 2025).

**Table 1.** Various measurements of species richness (α, β, γ). The direct measurement of species richness (γ_r_) is provided together with various estimates of extrapolated species richness, including the number of unobserved species. Chao’s (γ_c_), first order jackknife (γ_j_) and bootstrap estimators (γ_b_) are accompanied with standard errors. The second-order jackknife estimators are given in brackets. In the γ_r_ column, the number of species exclusive to one type of habitat or site type is shown in brackets. For instance, of the 51 species observed in dead wood, 17 were not found in either moss or soil samples. However, most of these species have been observed in only a very small number of samples.

| Diversity | | Alpha $\alpha$ | Beta $\beta$ | | Gamma $\gamma$ | | | |
| --- | --- | --- | --- | --- | --- | --- | --- | --- |
| | Group (number of samples) | Mean number of species per sample [range] | $\beta_r = \gamma_r / \alpha$ | $\beta_c = \gamma_c / \alpha$ | $\gamma_r$, total number of species (nb of species found only in one type of habitat or site) | $\gamma_c$, Chao's estimator | $\gamma_j$, First order Jackknife estimator (second order) | $\gamma_b$, Bootstrap estimator |
| **All samples together** | (89) | 5.3 [1–14] | 13.8 | 21 | 73 | 109± 19 | 100±7 (116) | 85±3.5 |
| **Microhabitats** | Moss (24) | 4.4 [1–8] | 7.5 | 12 | 33 (6) | 55±15 | 47±5 (57) | 39±2.5 |
|  | Soil (38) | 4.8 [1–11] | 9.8 | 14 | 47 (12) | 69±13 | 66±6 (77) | 56±3.2 |
|  | Wood (27) | 6.7 [1–14] | 7.6 | 11 | 51 (17) | 77±15 | 72±7 (85) | 60±3.8 |
| **Sites** | NPF (29) | 5.2 [1–10] | 7.7 | 11 | 40 (5) | 59±14 | 53±4 (62) | 46±2.5 |
|  | PAF (30) | 5.7 [2–14] | 8.9 | 13 | 51 (17) | 74±13 | 73±6 (85) | 61±3.4 |
|  | PNF (30) | 5 [1–12] | 9 | 15 | 45 (11) | 74±17 | 66±7 (79) | 54±4 |

We identified more species in 2022 (53) than in 2021 (43), but this may be due to the greater sampling effort (30 samples in 2021 and 59 in 2022). In fact, the species accumulation curves of the two years largely overlap (Figure 4A), indicating that the overall gamma diversity was roughly the same between the two years. Given this observation, and that our sampling protocol was not designed to allow a reliable comparison between years, we decided to group the two years together in the analysis.

Overall, we found slightly more species in the preserved anthropized forest (PAF, 51 species in total) than in the preserved natural forest (PNF, 45 species) and the non-preserved forest (NPF, 40 species, Table 1and Figure 4B). The species accumulation curves diverged progressively among the three sites (Figure 4B), showing that, despite the above-mentioned lack of significant differences on numbers of species per sample among sites (alpha diversity), the gamma diversity varied slightly among the sites. These variations between sites are reflected in the Jackknife and bootstrap estimates of gamma diversity, with higher diversity estimated in the PAF than in the NPF and PNF. The Chao estimator, meanwhile, finds the lowest diversity in the NPF (Table 1).

Regarding the microhabitats, the lower total number of species (gamma) was found in the moss samples (33), while about the same total species richness was found in the decaying wood (51) and in the forest soil and litter (47, Table 1 and Figure 3B). However, the species accumulation curves and the different estimates of gamma clearly show that, on average, dead wood is home to more springtail species than the other two microhabitats (Figure 4A), in line with what we observed for alpha diversity (Figure 3B).

Beta diversity is defined here as the ratio of gamma over alpha (β=γ/α) and can thus be interpreted as the effective number of distinct communities. Beta diversity varies between 7 and 21, depending on the scale at which it is measured and the method used (Table 1 and Figure 3B). The highest values are observed when all samples are grouped together, which makes sense, as at the scale of the whole community, beta diversity takes into account the spatial structure of communities between sites and between microhabitats. The maximum value of 21 indicates that an average of 21 distinct communities is needed to achieve the estimated total gamma diversity. At other spatial scales, beta diversity varies little between microhabitats and between sites (β_c_ ≍ ;11-15 & β_r_ ≍ 7-10), but still reaches maximum values for soil samples (β_c_ = 14) and for the PNF (β_c_ = 15, Figure 3B).

### Richness and Evenness

When analysing species evenness using Rényi diversity profiles, we found that the preserved natural forest site tends to exhibit higher diversity than the other two sites as evenness is progressively taken into account (Figure 5A). In terms of microhabitat types, diversity remained consistently higher in dead wood compared to soil or moss across the entire gradient of species evenness incorporation into the diversity measure (Figure 5B).

### Occupancy and Abundance

*Seira domestica* is the most abundant species across all sites studied, followed by *Folsomia peniculata*, a *Drepanura (sp.01), Heteromurus major, Ceratophysella stercoraria, Tomocerus minor, Orchesella cincta* and *Parisotoma notabilis* (Figure 6A). These species were dominant in terms of both total abundance and occupancy (ubiquity), appearing in more than about 30% of the samples (Figure 6B). For instance, *Drepanura sp.01* was present in about half of the samples.

Although on average global abundance increased with occupancy (Figure 7A), we found that this relationship was far from loglinear. In fact, we identified a group of relatively rare species, present in fewer than 10% of the samples for which there was no relationship between total abundance and occupancy. Some of these species were both rare and locally scarce (corresponding to “satellite” species), while others were rare but locally relatively abundant (corresponding to “urban” species like *Friesea sp.01*, *Folsomia sp.01*). In another group of species, the overall abundance increased with occupancy: from *Protaphorura bicampata* and *Mesaphorura yosii* to the ubiquitous and dominant *Seira domestica*, *Folsomia penicula* and *Drepanura sp.01* mentioned above (Figure 7A).

When considering abundance at a local scale (Figure 7B), a species – *Seira domestica* – stands out for being both globally ubiquitous and locally abundant. A core or ‘common species’ in other words. The other species form three main groups. The largest group includes species that are both rare and small in number when present (satellite or rare species). There are also species that are locally scarce, but present in a large proportion of the samples (rural species). Examples of these rural species include *Folsomia penicula*, *Tomocerus minor*, *Heteromurus major, Parisotoma notabilis..*. Finally, some species, such as *Proisotoma minuta*, *Pseudachorutes vasylii*, or *Schoettella ununguiculata* are infrequent but relatively abundant when present (“urban” species). Thus, due to this diversity in the relationship between occupancy and local abundance, the expected positive relationship between occupancy and local abundance is unclear in this community.

### Collembola Communities

The first two axes of the principal component analysis captured 31.8% of the total variation. On this projection plan, three groups of Collembola emerged (Figure 8A). The first group associates *Entomobrya nigrocincta* with *Seira domestica,* which may be related to the fact that *E. nigrocincta*, despite having much lower densities than *Seira domestica*, is almost always found in association with *S. domestica*. The second group includes *Tomocerus minor*, *Orchesella cincta*, *Heteromurus major* and the relatively locally abundant but less common *Entomobrya corticalis*. Finally, in the third group we find two Isotomidae (*Isotomiella minor* and *Folsomia penicula*), *Protaphorura bicampata* and *Mesaphorura yosii*. *Parisotoma notabilis* lies between the second and third group. Note that some rather common species such as *Drepanura sp.01*, *Ceratophysella stercoraria* contributed only marginally to these two main axes, which is consistent with the fact that the two main axes captured only slightly less than a third of the total variance.

**Figure 8.**
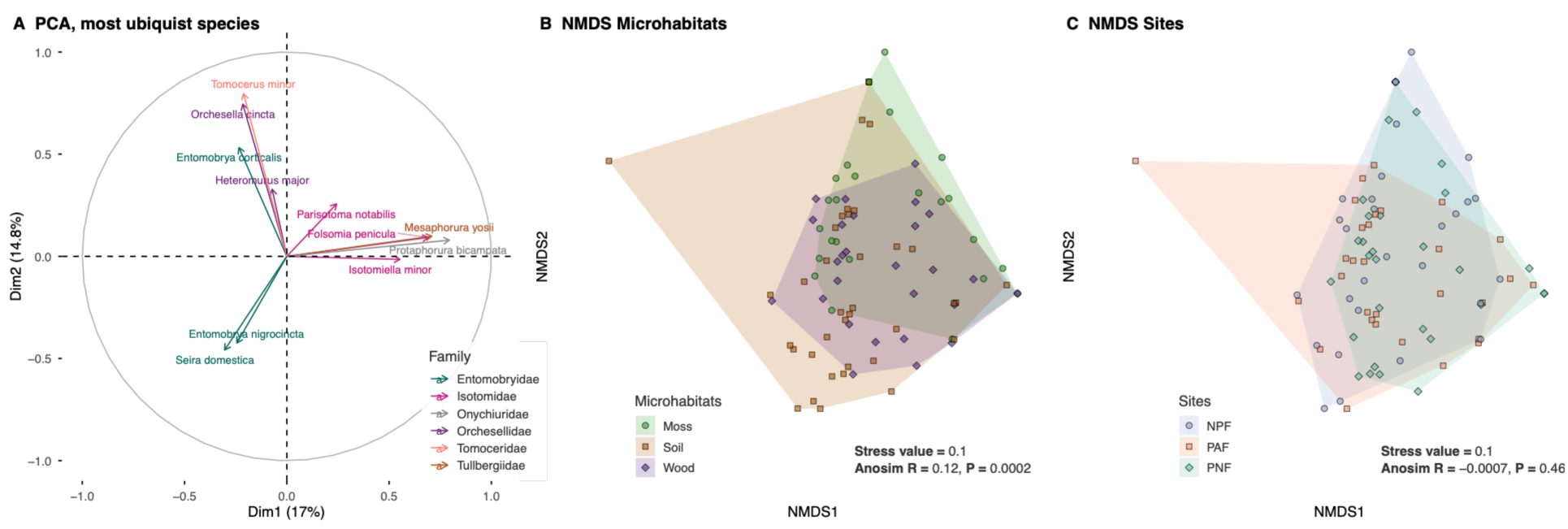
Principal Component Analysis of the most common species (A) and Non-Metric Multidimensional Scaling for the different microhabitats (B) of for the different sites (C). The different colours indicate the different families (A, cf. Figure 6) or the different micro-habitats (B) or sites (C).

When we plotted the samples on these two axes, we found very little community structuring between microhabitats or between sites (Figure not shown). NMDS and ANOSIM analyses indicated no clear structuring among sites (Figure 8C), but detected a significant, albeit weak, differentiation among microhabitats (Figure 8B, R = 0.117, P < 0.001), with dead wood communities showing the strongest separation.

None of the species was identified as an indicator species for any of the three sites (P > 0.03), but four species were identified as indicators of one or two microhabitats (Figure 9): *Folsomia peniculata* was associated with wood and soil microhabitats, *Orchesella cincta* with wood and moss microhabitats, *Isotomiella minor* was found almost exclusively in the soil, and *Lepidocytus sp.03* was mainly observed in dead wood.

**Figure 9.**
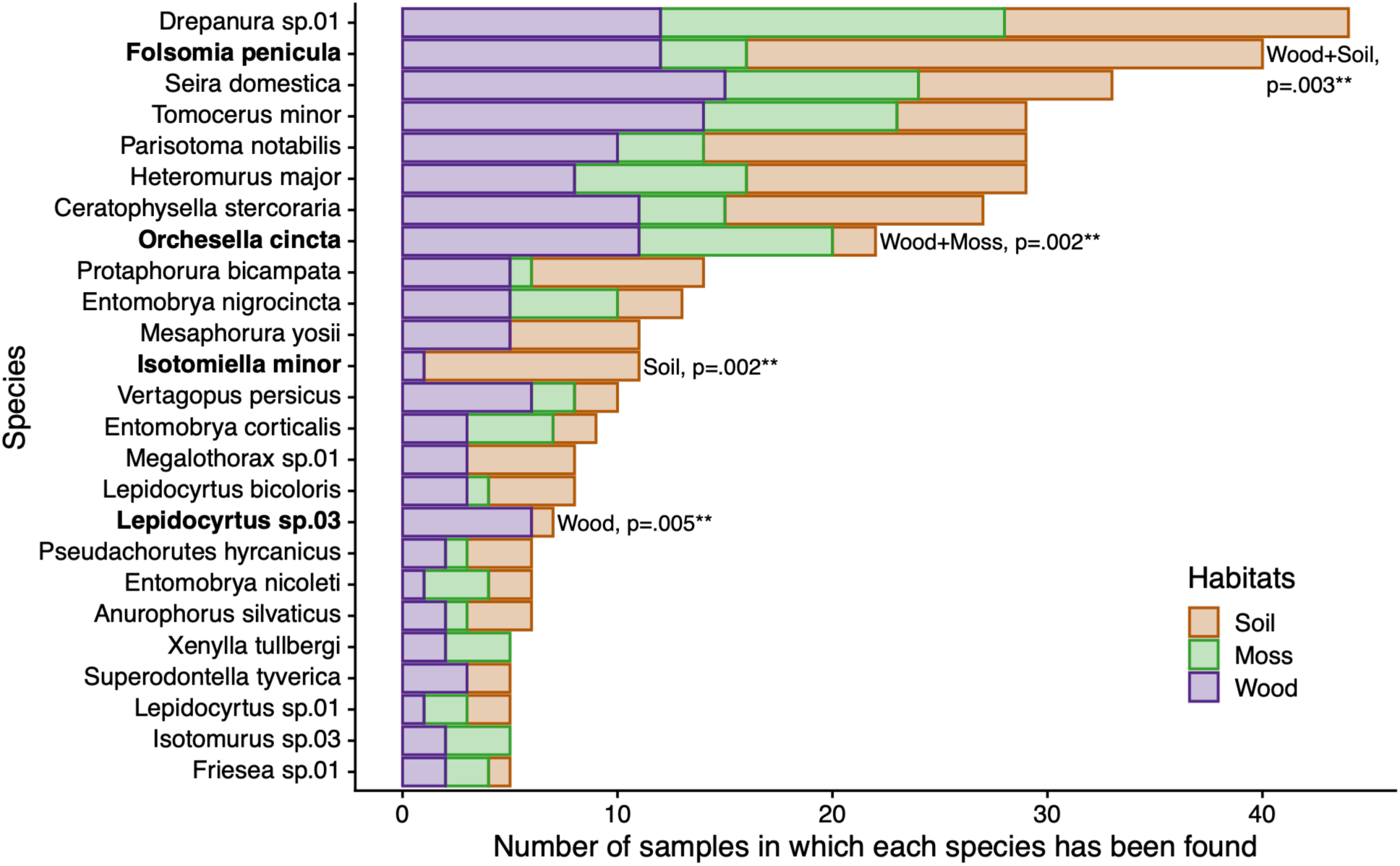
Distribution of the 25 species most frequently found in the three different types of microhabitats (observed in at least five samples). The species in bold are “indicator species”, i.e. those found to be significantly associated with one or two of the microhabitats. The corresponding p-values are listed next to the identified indicator species.

Excluding rare species (those present in fewer than 5 of 89 samples) leaves 25 species, of which 22 occurred at all three sites and 18 were present across all three microhabitats (Figure 9).

The distribution of species among the three microhabitats helps explain the weak microhabitat structuring detected in the previous analysis. Figure 9 shows that the ten most common species were present in all three microhabitats, whereas specialization to a single microhabitat was observed among only two less common species (*Isotomiella minor* and *Lepidocytus sp.03*).

To further explore the potential covariation of some species, as suggested by the three groups revealed in Figure 8A, we extended this data exploration with an analysis of co-occurrences between species (Figure 10). Of the 2628 species pair combinations, 234 pairs could be analysed, of which 205 (87.6%) revealed a random association between species, meaning that the observed number of co-occurrences for these pairs of species was not significantly different from the number of co-occurrences expected by chance (Griffith et al., 2016). In most cases, the presence of one species is not significantly dependent on the presence of another species. However, when co-occurrence does occur, it is predominantly driven by positive interactions. Of the 12.4% of non-random associations, we found 20 positive associations (8.5%) and 9 negative ones (3.8%, Figure 10A). For instance, *Tomocerus minor* is found more often than expected in association with *Orchesella cincta*, and *Folsomia penicula* is rarely observed in association with *Seira domestica* (7 co-occurrences compared to the 14 expected, Figure 10B).

**Figure 10.**
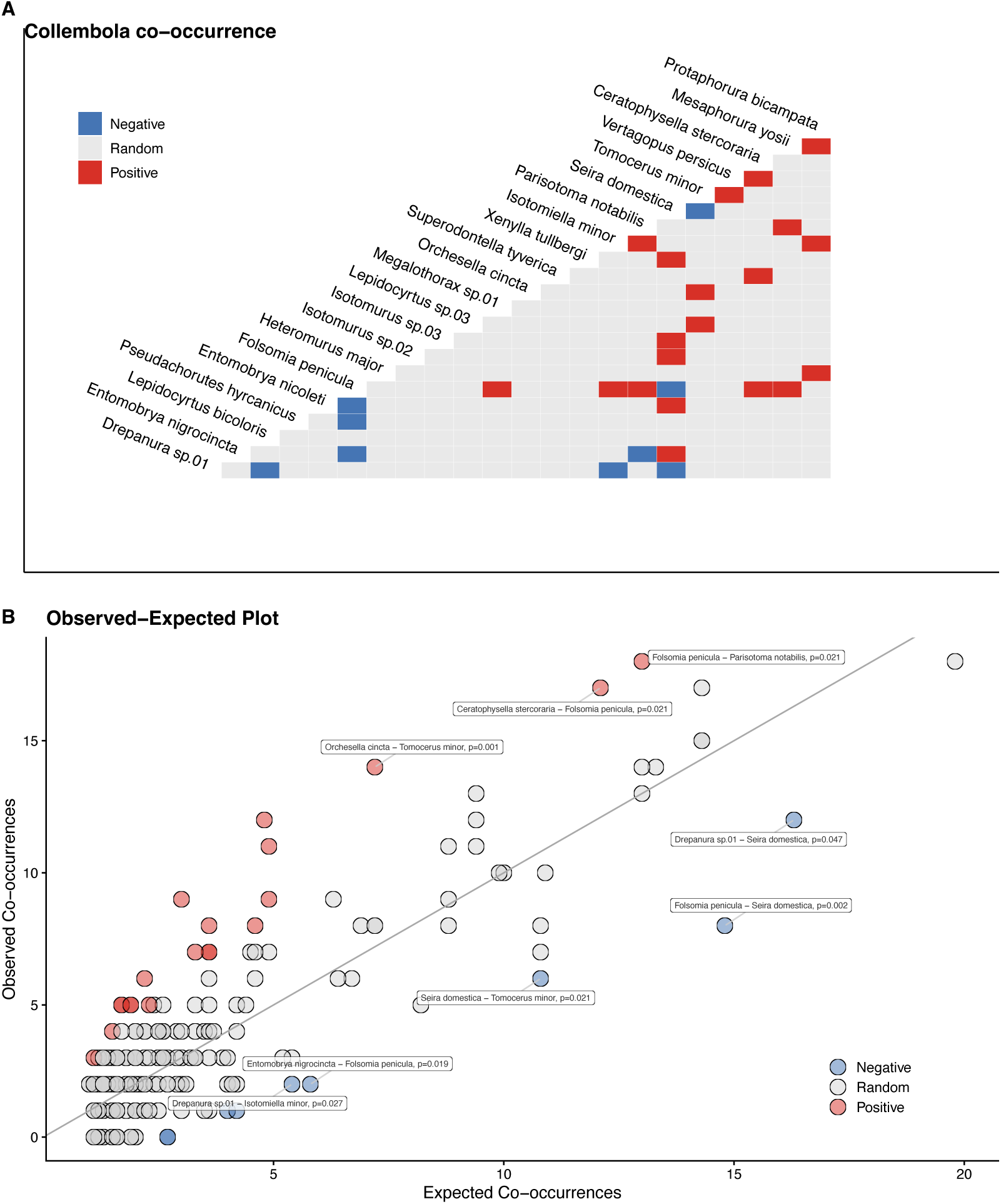
Co-occurrence between species. For most of the species’ pairs examined, no specific association was observed (random co-occurrence, grey). Species that are positively associated are underlined in red and those that are negatively associated are underlined in blue.

## Discussion

The Bula Forest, part of the ancient Hyrcanian forest and a UNESCO World Heritage Site, represents a rare example of old-growth forest with minimal human disturbance - particularly in its preserved natural sections. This study offers a comprehensive assessment of Collembola communities within this forest massif, focusing on key ecological attributes such as abundance, occupancy, diversity, richness, and evenness. Sampling took place close to the contact zone of two biodiversity hotspots, across three forest types with contrasting management histories (a preserved planted forest, a preserved natural forest and a non-preserved natural forest) and within three microhabitats in each site (wood, moss and soil/leaf litter).

### A particularly high level of species diversity

Despite the ecological importance of the Hyrcanian forests, little research has been devoted to the composition and structure of Collembola communities. Most previous studies have been limited to faunistic surveys. In the present study, 73 morphospecies were identified, spanning 39 genera and 12 families, with 49 species determined at the species level, highlighting a notably high level of diversity. For comparison, Mehrafrooz Mayvan et al. (2015a) previously reported 20 species across 16 genera and 9 families in the Semeskandeh Forest, including the description of a new endemic genus and species, *Persanura hyrcanica* (Mehrafrooz Mayvan et al., 2015b). In another study from Golestan National Park, 46 Collembola taxa were recorded (Khanahamdi et al., 2018). By 2019, a total of 107 species belonging to 14 families and 51 genera were known from the northern forests of Iran (Shayanmehr and Yahyapour, 2019), some of which, like *Pseudachorutes hyrcanicus*, are endemic (Shayanmehr et al., 2022).

Our sampling protocol enabled robust estimates of local diversity. Species richness per site or microhabitat ranged from 32 to 52 for observed diversity and from ∼40 to over 60 for estimated richness. There is also a marked difference between the gamma diversity and the local diversity (alpha), measured at the sample level, which averages around 5 to 6 species. This indicates a marked spatial structure in community composition (associated with a remarkably high beta diversity) and highlights the need to take numerous samples from the same site or type of microhabitat in order to obtain a reliable picture of the springtail species inhabiting these ecosystems.

Our gamma diversity measurements values are at the upper end of what is known for temperate forest Collembola communities, where average species richness typically hovers around 38, with 75% of studies reporting fewer than 51 species (Potapov et al., 2023). Overall, estimated total richness in the Bula Forest exceeded 90 species, which is remarkably high for a temperate ecosystem (Potapov et al., 2023; Addison et al., 2003; Sławska et al., 2017; Korboulewsky et al., 2021; Cassagne et al., 2006; Sławski and Sławska, 2020; Heiniger et al., 2014), but comparable to biodiversity levels observed in the central Pyrenees (Lauga-Reyrel and Lauga, 1995), some Canadian (Raymond-Léonard et al., 2020), Caucasus (Kuznetsova et al., 2019) or Chinese forests (Xie et al., 2025), although lower than in some relict forests in Russia (Kuznetsova et al., 2021) and Poland (Smolis and Skarżyński, 2003; Skarzynski and Smolis, 2006). However, methodological differences, particularly in sampling effort, site number, and taxonomic resolution, make direct comparisons difficult.

The Hyrcanian forests served as a critical refuge for temperate deciduous broad-leaved forests, particularly during the Quaternary glaciations (Browicz, 1989; Tohidifar et al., 2016). Their remarkable spatial and temporal stability likely contributed to the high diversity observed in springtail communities, as temporal stability is known to be a key driver of springtail diversity (Heiniger et al., 2014).

Our study expanded the scope beyond traditional soil and litter sampling by including mosses and dead wood, which significantly increased the number of observed species: approximately 55% more species were recorded thanks to these additional microhabitats (26 of the 73 species were found exclusively in moss or wood). This broader microhabitat coverage likely contributed to the higher diversity observed compared to other recent studies in Iran, such as those in the Zagros oak forests (Shayanmehr and Yahyapour, 2019; Ahmadi et al., 2023).

Nonetheless, our inventory is not exhaustive and some species may have been overlooked. Certain taxonomic groups, such as Symphypleona and Neelipleona remain underrepresented. Seasonal limitations – particularly the absence of autumn sampling – likely led to the omission of species with restricted temporal occurrence. Although our sampling effort had the advantage of being based on two years, which turned out to be relatively similar, it should be remembered that the composition of springtail communities can vary significantly from one year to the next, as shown by an 11-year monitoring of springtail communities in a pine forest in northern Latvia (Dirilgen et al., 2018). Longer term monitoring, over several seasons and years, across wider altitudinal gradients, different valleys and with deeper taxonomic resolution would be required to fully characterize the springtail communities of this forest massif.

### Spatial heterogeneity in Collembola species richness across microhabitats and forest sites

The cumulative species curves revealed notable differences in species richness both between microhabitats and across different sites (Figure 4C). The divergence of the cumulative species curves between the three microhabitats, and between the three sites, highlight spatial heterogeneity in diversity at two distinct spatial scales.

Across sites, our study demonstrates that Collembola gamma diversity is highest in dead wood and lowest in mossy microhabitats. This difference is also evident, albeit to a lesser extent, at the sample level, with a slightly higher alpha diversity in the deadwood samples (Figure 3B). Moreover, this difference between microhabitats persisted, and even intensified, when the relative abundance of each species was considered (Figure 5B): dead wood not only supports a slightly greater number of Collembola species but also fosters communities with more balanced relative abundances. Similar findings have been reported in Quebec forests, where 74 Collembola species were documented in dead wood microhabitats (Raymond-Léonard et al., 2020). However, in a couple of Netherland forests, Collembola species diversity (29 species in total) was found to be similar between moss, soil and dead wood, but the Shannon or Simpson diversity index was highest for soil, followed by dead wood and finally moss (Fujii et al., 2023).

Our results reinforce the view that dead wood in forests serves as a critical habitat for a high diversity of springtails, as well as other taxonomic groups (Skubała and Marzec, 2013). This underscores the importance of conserving various types of dead wood in forest ecosystems, as they provide rich and diverse microhabitats (Seibold et al., 2016) and contribute significantly to habitat heterogeneity (Fujii et al., 2023).

Furthermore, our study shows that the specific diversity of Collembola communities tends to differ between the three sites. The site with the highest species richness is the preserved anthropogenic forest, whereas the non-preserved forest exhibits the lowest species richness. But this difference between sites is only apparent when looking at cumulative species curves and gamma diversity, as mean alpha diversity is found to be similar across sites. This suggests that the observed variation in diversity among the forest patches is driven less by differences in local species richness than by differences in the spatial structuring of communities at metre to decimetre scales within each site. Such patterns may reflect greater spatial heterogeneity in the most species-rich sites.

Interestingly, when considering not only species numbers but also their relative abundance, the preserved natural forest emerges as the most diverse in terms of Collembola communities (Figure 5A), with this site’s diversity exceeding that of the preserved anthropized forest, despite the fact that the latter had the highest number of species recorded (the Rényi diversity profiles intersect). This means that the Collembola communities in the preserved natural forest are less dominated by dominant species than the other sites. In communities dominated by one or a few highly abundant springtails, competitive exclusion may intensify, reducing the populations of less competitive species (Hishi et al., 2022). This process can amplify demographic disparities within the community, leading to skewed species abundances. Our observations support the importance of preserving natural forests in order to maintain communities that are both species-rich and balanced in terms of relative abundance, aligning with previous studies (Deharveng, 1996; Oliveira and Deharveng, 1995; Susanti et al., 2021).

Nevertheless, this interpretation should be treated with caution, as our study included only a single site for each forest management type, which limits our ability to attribute the observed differences to management alone. Other factors such as altitude and local environmental conditions also differ between sites and may contribute to the patterns observed.

Stepping back to a broader perspective, comparisons between the Bula Forest natural area, which is highly protected (UNESCO World Heritage), and less protected Hyrcanian forests that experience ongoing disturbances (e.g. logging, land-use change, road construction, recreational infrastructure) suggest a possible effect of protection status on Collembola communities. In the well-protected Bula Forest, Collembola diversity and density appear higher, with 47 species and on average 12,400 individuals per square metre in soil samples, compared to about 7,600 ind. m⁻² reported for the less-protected Semeskaneh Forest (Mehrafrooz Mayvan et al., 2015a). While these comparisons are not strictly controlled and should be interpreted as indicative rather than causal, they are consistent with the hypothesis that strong conservation measures may help maintain elevated levels of Collembola biodiversity in these forest ecosystems.

### Abundance and occupancy varied a lot between species

*Seira domestica* was the most abundant species, both locally and globally, and one of the most frequently observed in our different samples in the Bula Forest. This species therefore has the characteristics of a common - or dominant, or core - species within our study site although it has been observed in less than 50% of the samples (Avolio et al., 2019; Collins et al., 1993). It is interesting to note that it is the only one that is common in these communities, while in Semeskandeh Forest (Mehrafrooz Mayvan et al., 2015a; Ghasemi Charati et al., 2021) and in three other Hyrcanian forests, namely Zare, Dohezar and Langar (Yahyapour et al., 2019), *Heteromurus major* was the most abundant species. However, apart from the total number of individuals, which can vary widely due to different population dynamics, our study also found that *Heteromurus major* was one of the most ubiquitous species in our samples.

Despite a clear positive relationship between occupancy and global abundance, the expected positive relationship between local abundance and occupancy was not supported in our study because the collembola community was not composed only of core and satellite species, but also of rural and urban species. More precisely we found one core species (*S. domestica*) and numerous satellite species. The community was also composed of nine urban species exhibiting low occupancy yet high local abundance when present (such as *Proisotoma minuta* and *Pseudachorutes vasylii)* and seven rural one, largely distributed but in small numbers when present. This pattern raises critical questions about the ecological and demographic processes underlying such distributions. It is interesting to find this type of species distribution within animal communities among springtails, similar to what has been observed, for example, in *Drosophila* successions on fallen fruit (Deconninck et al., 2024). Identifying three types of rare species within our communities (satellite, rural, or urban species) raises the question of the mechanisms underlying these differences; these may be linked to variations in growth, colonization, dispersal, and extinction rates, as well as to specific ecological preferences. For instance, urban species may be characterized by high growth rates and limited dispersal abilities that may restrict these species to localized patches where they proliferate (Hanski, 1991). Further studies are required to determine whether these findings hold across broader spatial or taxonomic contexts, as well as to investigate potential mechanisms that could be linked to functional traits (Raymond-Léonard et al., 2018; Joimel et al., 2024), unexplored niche dimensions – such as microclimatic preferences or resource specialization – that were not captured in our analyses, and the potential influence of interspecific interactions like competition. Additionally, temporal fluctuations in population sizes, which are inherently difficult to detect in cross-sectional studies, may play a key role in shaping these patterns (Pozo et al., 1986; Kuznetsova, 2007; Mehrafrooz Mayvan et al., 2015a). Clarifying these mechanisms will help distinguish whether the observed structuration of the Collembola communities into four unbalanced groups in the occupancy-abundance arise from species-specific traits, unmeasured ecological constraints, or broader community-level processes governing springtail assemblages. Finally, one might wonder whether the high gamma biodiversity could be linked to the near-total absence of core species in these communities, thereby allowing for a high abundance of satellite species, and to a lesser extend urban or rural species. This large number of rare species should also be considered in relation to the high beta-diversity values measured.

### Differences between families

Comparing the abundance and occupancy of springtail species reveals that most families contain a few dominant species (numerous and widespread) and others species that are much rarer and less abundant. Notably, Figure 6 shows that dominant species belong to five different families, with Neanuridae standing out in our dataset in that it includes species that are never frequent, but sometimes locally abundant. This pattern could be due to strong spatial population structure in the form of dense local aggregations combined with limited dispersal or reduced mobility. Neanuridae lack a functional furca and exhibit particularly slow locomotion compared to other springtails (Paśnik and Smolis, 2024; Ponge et al., 2006). Their spatial distribution is thus probably largely shaped by fine-scale microhabitat heterogeneity (organic matter patches, soil aggregate, wood decomposition gradients) characteristic of forest soils (Berg, 2012). Additionally, their scratching/piercing mouthparts and scavenger/carnivorous diet (Malcicka et al., 2017) may further constrain habitat suitability, potentially explaining their patchy distribution. Further ecological studies are needed to test these hypotheses through detailed observations of their microhabitat preferences and movement ecology. A natural extension of this study would be to adopt a functional trait-based approach to examine how microhabitats and sites structure collembola community functional diversity, rather than taxonomic diversity alone (Reis et al., 2016; Joimel et al., 2024; Potapov et al., 2020b; Holmstrup et al., 2018; Reis et al., 2016).

### Most common species are generalists

Contrary to findings reported in other studies, particularly in forests in the Netherlands, where the specific compositions of springtail communities in dead wood differ from those in mosses or the soil (Fujii et al., 2023), we were unable to identify any marked differences in community composition between microhabitats (or between sites). This is linked to the fact that most species do not have very marked preferences for any of the three microhabitats (or sites). Indeed, the most common species are generalists in the sense that they can colonize all three types of microhabitats. Some of these species are composed of quite large and mobile specimens (*Depanura sp.*, *S. domestica*, *H. major*, *T. minor*), and it is possible that the territory explored by these individuals may encompass the different types of microhabitats surveyed. But this is questionable for other smaller species such as *Parisotoma notabilis*.

We found only four indicator species, i.e. species showing a marked preference for or avoidance of one of the microhabitats. *Isotomiella minor* showed a strong preference for soil, as previously reported in Dutch forests for example (Fujii et al., 2023). However, *Orchesella cincta*’s association with dead wood and moss was not evident in the Dutch forests, and *Parisotoma notabilis* was not identified as an indicator species in our study, although it was clearly associated with soil habitats in Dutch forests (Fujii et al., 2023).

### Co-occurrence analysis reveals the existence of covariation between certain pairs of species

Our co-occurrence analysis shows that, for most species’ pairs, the interactions are neutral, meaning the presence of one species is not influenced by the presence of another. This coexistence is probably driven by strong horizontal and vertical spatial heterogeneity in these habitats, which generate diverse resource availability and fine-scale microclimatic gradients. These factors create a mosaic of microhabitats that promotes niche partitioning and supports high species coexistence (Berg, 2012). We were also able to demonstrate the presence of several positive associations between species, as well as a smaller number of negative associations. The latter could be a sign of interspecific competitive interactions. However, the origin and ecological significance of both positive and negative associations, some of which could be ‘interactions’, remain to be elucidated, as interspecific relations among springtails are not well understood. It is known that intraspecific competition can be complex, combining resource exploitation and interference between age classes (Mallard et al., 2019; Le Bourlot et al., 2014). Interspecific interactions between Collembola can also be complex and sometimes asymmetrical (+/-) (Theenhaus et al., 1999), and may further depend on temperature and on the presence of predators (Thakur et al., 2017). Beyond attraction followed by competition for shared resources, predation between Collembola can occur (in particular through oophagy), and potential negative density-dependent effects can occur. Laboratory experiments could improve our understanding of the conditions and mechanisms by which the abundance of one species can be increased with the presence of another. To our knowledge, this type of analysis has rarely been carried out on Collembola. It would be interesting to see if the same type of associations between species could be identified in other datasets and forest ecosystems. Our findings could also help to select species pairs for experimental studies, helping us to better understand the ecological causes of these associations (e.g. shared preferences, competition, exploitation or mutualism).

## Conclusion

Despite being limited to just three sites within a relatively small portion of the Hyrcanian Forest, and acknowledging that some Collembola groups remain understudied, our study reveals exceptionally high beta and gamma diversity. This richness, observed in relict forests at the intersection of two major biodiversity hotspots, underscores the high ecological value of the region, further recognized by its inclusion on the UNESCO World Heritage List. Our findings also highlight the critical importance of conserving protected forests, particularly those with abundant deadwood, which provides spatial structure and essential microhabitats that support both species-rich and balanced Collembola communities.

Contrary to initial expectations, we found no strong structuring of Collembola communities between sites or microhabitats. Instead, these communities appear to be dominated by generalist species, with only a minority associated with one or two specific microhabitats. However, we did observe marked differences between species, and even among certain families, in abundance and occupancy, likely driven by fine-scale ecological and demographic variation. While Collembola communities largely seem to consist of independently co-occurring species, our analysis also revealed partial structuring due to both positive and negative interspecific associations.

These findings open new avenues for research into the direct and indirect interspecific interactions shaping Collembola assemblages in forest ecosystems. They also emphasize the need to preserve forest structural complexity, particularly deadwood, to maintain the diversity of these often-overlooked communities.

## Authors Contributions

TT, EYL, CDH and MS conceived the ideas and designed the sampling protocol; EYL took the samples in the field, set up the Berlese equipment and sorted the animals. MGS did the soil analysis. MS, CD and EYL identified the specimens. TT and CDH organized the database. TT did the statistical analysis, produced the graphics and wrote the manuscript. All authors gave final approval for publication.

## Acknowledgments

We would like to thank the Centre for International Scientific Studies & Collaboration (CISSC), Ministry of Science Research and Technology (MSRT, national ID: 14002786851) and Deputy of Research and Technology of Sari University of Agricultural Sciences and Natural Resources (SANRU, National ID: 14002820777) for providing appropriate condition to conduct the research (National ID: 14002820777). We also would like to thank our contacts from Campus France’s Partenariat Hubert Curien partnership and from the French Embassy in Teheran for the trust they have placed in our collaborative project and for their financial and logistical support. MS also would like to thank Mr. Hadrian Rozier (Associate of Scientific and Academic Cooperation) to accelerate her stay in Paris. We are thankful to Olivier Blight, Olivier Chabrerie and Tania de Almeida for their valuable comments and suggestions.

## Data & Script

Data is available here: https://zenodo.org/records/15658167

The script written to handle the data, run the statistical analysis and produce the figures is here: https://zenodo.org/records/15658467

## References

Addison, J A, J A Trofymow, and V G Marshall (2003), ‘Abundance, species diversity, and community structure of Collembola in successional coastal temperate forests on Vancouver Island, Canada’, Applied Soil Ecology, 24 (3), 233–46.

Ahmadi, Somayeh, et al. (2023), ‘Addition to Iranian Springtails fauna and a checklist of the Collembola from Kurdistan province’, Journal of Insect Biodiversity and Systematics, 9 (1), 1–16.

Akhani, Hossein, et al. (2010), ‘Plant biodiversity of Hyrcanian relict forests, N Iran: an overview of the flora, vegetation, palaeoecology and conservation’, Pakistan Journal of Botany, 42 (1), 231–58.

Avolio, Meghan L, et al. (2019), ‘Demystifying dominant species’, New Phytol, 223 (3), 1106–26.

Bellinger, P.F., K.A. Christiansen, and F. Janssens (2024), ‘Checklist of the Collembola of the World’,

Berg, Matty P (2012), ‘Patterns of biodiversity at fine and small spatial scales’, Soil ecology and ecosystem services (136–52.

Bhagawati, Sudhansu, et al. (2021), ‘Diversity of Soil Dwelling Collembola in a Forest, Vegetable and Tea Ecosystems of Assam, India’, Sustainability, 13 (22), 12628.

Blackburn, Tim M., Phillip Cassey, and Kevin J. Gaston (2006), ‘Variations on a Theme: Sources of Heterogeneity in the Form of the Interspecific Relationship between Abundance and Distribution’, Journal of Animal Ecology, 75 (6), 1426–39.

Browicz, Kazimierz (1989), ‘Chorology of the Euxinian and Hyrcanian element in the woody flora of Asia’, Plant systematics and Evolution, 162 (1), 305–14.

Brown, James H. (1984), ‘On the Relationship between Abundance and Distribution of Species’, The American Naturalist, 124 (2), 255–79.

Caruso, Tancredi, et al. (2013), ‘Biotic interactions as a structuring force in soil communities: evidence from the micro-arthropods of an Antarctic moss model system’, Oecologia, 172 (2), 495–503.

Cassagne, Nathalie, et al. (2006), ‘Endemic Collembola, privileged bioindicators of forest management’, Pedobiologia, 50 (2), 127–34.

Chahartaghi, Masoumeh, et al. (2005), ‘Feeding guilds in Collembola based on nitrogen stable isotope ratios’, Soil Biology and Biochemistry, 37 (9), 1718–25.

Collins, Scott L, Susan M Glenn, and David W Roberts (1993), ‘The hierarchical continuum concept’, J Vegetation Science, 4 (2), 149–56.

D’Haese, Cyrille, et al. (2025), ‘An Initial Assessment of the Collembola (Hexapoda) Fauna of the Bula Hyrcanian Forest (Iran) at the Interface of Two Biodiversity Hotspots’, Soil Organims, 97

De Cáceres, Miquel and Pierre Legendre (2009), ‘Associations between species and groups of sites: indices and statistical inference’, Ecology, 90 (12), 3566–74.

Deconninck, Gwenaëlle, et al. (2024), ‘Fallen fruit: A backup resource during winter shaping fruit fly communities’, Agri and Forest Entomology, 26 (2), 232–48.

Deharveng, Louis (1996), ‘Soil Collembola Diversity, Endemism, and Reforestation: A Case Study in the Pyrenees (France)’, Conservation Biology, 10 (1), 74–84.

Dirilgen, Tara, et al. (2018), ‘Analysis of spatial patterns informs community assembly and sampling requirements for Collembola in forest soils’, Acta Oecologica, 86 23–30.

Faraway, Julian J. (2016), Extending the Linear Model with R: Generalized Linear, Mixed Effects and Nonparametric Regression Models, (Chapman & Hall/CRC Texts in Statistical Science) 413.

Filser, Juliane (2002), ‘The role of Collembola in carbon and nitrogen cycling in soil’, Pedobiologia, 46 (3-4), 234–45.

Fjellberg, Arne (1998), The Collembola of Fennoscandia and Denmark, (Leiden; Boston: Brill).

Fjellberg, Arne (2007), The Collembola of Fennoscandia and Denmark, Part II: Entomobryomorpha and Symphypleona, (BRILL) 272.

Fujii, Saori, et al. (2023), ‘Downed deadwood habitat heterogeneity drives trophic niche diversity of soil-dwelling animals’, Soil Biology and Biochemistry, 187 109193.

Gange, A (2000), ‘Arbuscular mycorrhizal fungi, Collembola and plant growth’, Trends in Ecology & Evolution, 15 (9), 369–72.

Gaston, Kevin J. (1996), ‘The Multiple Forms of the Interspecific Abundance-Distribution Relationship’, Oikos, 76 (2), 211–20.

Gaston, Kevin J., Tim M. Blackburn, and John H. Lawton (1997), ‘Interspecific Abundance-Range Size Relationships: An Appraisal of Mechanisms’, Journal of Animal Ecology, 66 (4), 579–601.

Gee, Glendon W and James W Bauder (1986), ‘Particle-size analysis’, in Klute, Arnold (ed.), Methods of soil analysis: Part 1. Physical and mineralogical methods (Agronomy Monograph No. 9, Madison, Wisconsin: Soil Science Society of America and the American Society of Agronomy), 383–411.

Ghasemi Charati, Mahdiyeh, et al. (2021), ‘Introduction to class of Collembola as soil mesofauan from Semeskandeh mixed forest (Hyrcanian region)’, Ecology of Iranian Forest, 9 (18), 115–26.

Griffith, Daniel M., Joseph A. Veech, and Charles J. Marsh (2016), ‘cooccur: Probabilistic Species Co-Occurrence Analysis in R’, *Journal of Statistical Software*, Code Snippets, 69 (2), 1–17.

Haghdoost, Niloufar, et al. (2011), ‘Conversion of Hyrcanian degraded forests to plantations: Effects on soil C and N stocks’, *Ann*. Biol. Res, 2 385–99.

Hanski, Ilkka (1991), ‘Single-species metapopulation dynamics: concepts, models and observations’, Biological Journal of the Linnean Society, 42 (1-2), 17–38.

Harta, István, et al. (2021), ‘Collembola communities and soil conditions in forest plantations established in an intensively managed agricultural area’, Journal of Forestry Research, 32 (5), 1819–32.

Heiniger, Charlène, et al. (2014), ‘Effect of habitat spatiotemporal structure on collembolan diversity’, Pedobiologia, 57 (2), 103–17.

Hishi, Takuo, Erika Kawakami, and Ayumi Katayama (2022), ‘Changes in the abundance and species diversity of Collembola community along with dwarf bamboo density gradient in a mountainous temperate forest of Japan’, Applied Soil Ecology, 180 104606.

Hoffman, M, et al. (2016), ‘Biodiversity hotspots (version 2016.1)[Data set]’, Zenodo,

Holmstrup, Martin, et al. (2018), ‘Functional diversity of Collembola is reduced in soils subjected to short-term, but not long-term, geothermal warming’, Functional Ecology, 32 (5), 1304–16.

Homami Totmaj, Leila, et al. (2021), ‘Late Holocene Hyrcanian forest and environmental dynamics in the mid-elevated highland of the Alborz Mountains, northern Iran’, Review of Palaeobotany and Palynology, 295 104507.

Hopkin, Stephen P (1997), Biology of the Springtails (Insecta: Collembola), (Oxford: Oxford University Press).

Joimel, Sophie, et al. (2024), ‘Trait concepts, categories, and databases in soil invertebrates ecology– ordering the mess’, Soil organisms, 96 (3), 151–56. (ed.) (2012), *Capbryinae & Entomobryini*, Synopses on Palaeartic Collembola, 7/1; Senckenberg: Museum of Natural History Görlitz) 390.

Jost, Lou (2007), ‘Partitioning diversity into independent alpha and beta components’, Ecology, 88 (10), 2427–39.

Kassambara, Alboukadel (2017), Practical Guide To Principal Component Methods in R, (STHDA) 169.

Khanahamdi, Susan, Masoumeh Shayanmehr, and Mohammad Ali Bahmanyar (2018), ‘New record of Friesea afurcata Tullberg, 1869 (Collembola, Neanuridae) in Golestan national Park (Hyrcanian forests), Iran’, Journal of Insect Biodiversity and Systematics, 4 (3), 141–146-141.

Kindt, Roeland and Richard Coe (2005), Tree diversity analysis: a manual and software for common statistical methods for ecological and biodiversity studies, (World Agroforestry Centre).

Koleff, Patricia, Kevin J. Gaston, and Jack J. Lennon (2003), ‘Measuring beta diversity for presence–absence data’, Journal of Animal Ecology, 72 (3), 367–82.

Korboulewsky, N., et al. (2021), ‘Effect of tree mixture on Collembola diversity and community structure in temperate broadleaf and coniferous forests’, Forest Ecology and Management, 482 118876.

Kuznetsova, N and N Ivanova (2020), ‘Diversity of Collembola under various types of anthropogenic load on ecosystems of European part of Russia’, Biodivers Data J, 8 e58951.

Kuznetsova, N, et al. (2021), ‘The extremely high diversity of Collembola in relict forests of Primorskii Krai of Russia’, Biodivers Data J, 9 e76007.

Kuznetsova, N. A., et al. (2019), ‘Structure of the Species Diversity of Soil Springtails (Hexapoda, Collembola) in Pine Forests of the Caucasus and the Russian Plain: a Multi-Scale Approach’, Entomological Review, 99 (2), 143–57.

Kuznetsova, NA (2007), ‘The long-term dynamics of collembola populations in forest and meadow ecosystems’, Entomological Review, 87 (1), 11–24.

Lauga-Reyrel, F and J Lauga (1995), ‘Collembola of cold Pyrenean habitats’, Hydrobiologia, 312 59–74.

Le Bourlot, Vincent, Thomas Tully, and David Claessen (2014), ‘Interference versus exploitative competition in the regulation of size-structured populations’, The American Naturalist, 184 (5), 609–23.

Lê, Sébastien, Julie Josse, and François Husson (2008), ‘FactoMineR: An R Package for Multivariate Analysis’, Journal of Statistical Software, 25 (1), 1–18.

Lindberg, Niklas, Jan B. Engtsson, and Tryggve Persson (2002), ‘Effects of experimental irrigation and drought on the composition and diversity of soil fauna in a coniferous stand’, Journal of Applied Ecology, 39 (6), 924–36.

Loranger, Gladys, et al. (2001), ‘Does soil acidity explain altitudinal sequences in collembolan communities?’, Soil Biology and Biochemistry, 33 (3), 381–93.

Loreau, Michel, Otto Kinne, and Anders Pape Møller (2010), The challenges of biodiversity science, (17; International Ecology Institute Oldendorf/Luhe, Germany).

Malcicka, Miriama, Matty P. Berg, and Jacintha Ellers (2017), ‘Ecomorphological adaptations in Collembola in relation to feeding strategies and microhabitat’, European Journal of Soil Biology, 78 82–91.

Mallard, François, et al. (2019), ‘From individuals to populations: How intraspecific competition shapes thermal reaction norms’, Functional Ecology, 34 669–83.

Marcon, Eric (2015), ‘Mesures de la biodiversité’, (AgroParisTech).

Mari-Mutt, JA (1979), ‘A Revision of the Genus Dicranocentrus Schött (Insecta: Collembola: Entomobryidae)’, Bulletin of the University of Puerto Rico, 259 1–79.

Mehrafrooz Mayvan, Mahmood, Masoumeh Shayanmehr, and Stefan Scheu (2015a), ‘Depth distribution and inter-annual fluctuations in density and diversity of Collembola in an Iranian Hyrcanian forest: Additional contribution’, Soil Organisms, 87 (3), 239–247-239.

Mehrafrooz Mayvan, Mahmood, et al. (2015b), ‘*Persanura hyrcanica*, a new genus and species of Neanurinae (Collembola: Neanuridae) from Iran, with a key to genera of the tribe Neanurini’, Zootaxa, 3918 (4), 552–552.

Mehrafrooz Mayvan, Mahmood, et al. (2022), ‘Density, diversity, and seasonal fluctuations in soil Collembola in three differently managed ecosystems in North Khorasan, Iran’, Turkish Journal of Zoology, 46 (1), 115–28.

Nelson, D. W. and L. E. Sommers (1996), ‘Total Carbon, Organic Carbon, and Organic Matter’, in Sparks, D.L., et al. (eds.), Methods of Soil Analysis. Part 3. Chemical Methods (SSSA Book Series No. 5, Madison, Wisconsin: Soil Science Society of America and the American Society of Agronomy), 961–1010.

Noroozi, Jalil, et al. (2019), ‘Endemic diversity and distribution of the Iranian vascular flora across phytogeographical regions, biodiversity hotspots and areas of endemism’, Sci Rep, 9 (1), 12991.

Noroozi, Jalil, et al. (2023), ‘Hotspots of (sub)alpine plants in the Irano-Anatolian global biodiversity hotspot are insufficiently protected’, Diversity and Distributions, 29 (2), 244–53.

Nourzad Moghaddam, Mohsen, et al. (2018), ‘Determining basic concepts to formulate the Hyrcanian forest policy’, Iranian Journal of Forest, 10 (3), 373–87.

Oksanen, J, et al. (2026), vegan: Community Ecology Package, (R package version 2.7-3, The R Foundation).

Oliveira, EP and L Deharveng (1995), ‘Response of soil Collembola (Insecta) communities to forest disturbance in central Amazonia (Brazil)’, Functioning and Dynamics of Natural and Perturbed Ecosystems. Technique et Documentation, Lavoisier, Intercept Ltd, 361–76.

Paśnik, Grzegorz and Adrian Smolis (2024), ‘Is the Current Systematic Subdivision of the Subfamily Neanurinae (Collembola, Neanuridae) Still Valid? Testing the Monophyly and Phylogenetic Relationships of Currently Established Tribes of the Neanurinae’, Insects, 15 (9), 672.

Paul, D., A. Nongmaithem, and L. K. Jha (2011), ‘Collembolan Density and Diversity in a Forest and an Agroecosystem’, Open Journal of Soil Science, 01 (02), 54–60.

Petersen, H (2011), ‘Collembolan communities in shrublands along climatic gradients in Europe and the effect of experimental warming and drought on population density, biomass and …’, Soil Organisms,

Petersen, Henning (2002), ‘General aspects of collembolan ecology at the turn of the millennium’, Pedobiologia, 46 246–60.

Ponge, J.F. and S. Salmon (2013), ‘Spatial and taxonomic correlates of species and species trait assemblages in soil invertebrate communities’, Pedobiologia, 56 (3), 129–36.

Ponge, JF, et al. (2006), ‘Decreased biodiversity in soil springtail communities: the importance of dispersal and landuse history in heterogeneous landscapes’, Soil Biology & Biochemistry, 38 (5), 1158–61.

Potapov, AM, et al. (2023), ‘Globally invariant metabolism but density-diversity mismatch in springtails’, Nat Commun, 14 (1), 674.

Potapov, AM, et al. (2024), ‘Global fine-resolution data on springtail abundance and community structure’, Sci Data, 11 (1), 22.

Potapov, Anton, et al. (2020a), ‘Towards a global synthesis of Collembola knowledge: challenges and potential solutions’, soil organisms, 92 (3), 161–88.

Potapov, Anton, et al. (2020b), ‘Towards a global synthesis of Collembola knowledge–challenges and potential solutions’, Soil organisms, 92 (3), 161–88.

Dunger, Wolfram (ed.) (2001), Isotomidae, 3; Staatliches Museum für Naturkunde Görlitz)

Pozo, J, D Selga, and JC Simon (1986), ‘Studies on the collembolan populations of several plant communities of the Basque Country (Spain)[diversity, vertical distribution; forests (pine, oak, beech), meadow]’, Revue d’Ecologie et de Biologie du Sol (France), 23 (2),

Raymond-Léonard, Laura J., Mathieu Bouchard, and I. Tanya Handa (2020), ‘Dead wood provides habitat for springtails across a latitudinal gradient of forests in Quebec, Canada’, Forest Ecology and Management, 472 118237.

Raymond-Léonard, LJ, et al. (2018), ‘Springtail community structure is influenced by functional traits but not biogeographic origin of leaf litter in soils of novel forest ecosystems’, Proc Biol Sci, 285 (1879), 20180647.

Reis, Filipa, et al. (2016), ‘The use of a functional approach as surrogate of Collembola species richness in European perennial crops and forests’, Ecological Indicators, 61 676–82.

Rényi, Alfréd (1961), ‘On measures of entropy and information’, Proceedings of the Fourth Berkeley Symposium on Mathematical Statistics and Probability, Volume 1: Contributions to the Theory of Statistics 547–61.

Rhoades, JD (1982), ‘Soluble salts’, in Page, Albert Lee, R. H. Miller, and D. R. Keeney (eds.), Methods of soil analysis: Part 2. Chemical and microbiological properties (Agronomy Monograph 9, Madison, Wisconsin: Soil Science Society of America), 167–79.

Roberts, David W (2023), ‘labdsv: Ordination and multivariate analysis for ecology’, R package version 2.1.0, 1 (1),

Rusek, Josef (1998), ‘Biodiversity of Collembola and their functional role in the ecosystem’, Biodiversity and Conservation, 7 (9), 1207–19.

Saeei, K. (1950), Silviculture, (2; University of Teheran) 195.

Sagheb Talebi, Khosro, Toktam Sajedi, and Mehdi Pourhashemi (2014), Forests of Iran. A Treasure from the Past, a Hope for the Future, (Dordrecht: Springer Netherlands) 271.

Seibold, Sebastian, et al. (2016), ‘Dead-wood addition promotes non-saproxylic epigeal arthropods but effects are mediated by canopy openness’, Biological Conservation, 204 181–88.

Shayanmehr, M, et al. (2013), ‘An introduction to Iranian Collembola (Hexapoda): an update to the species list’, Zookeys, (335), 69–83.

Shayanmehr, M, et al. (2022), ‘New Pseudachorutes species Tullberg, 1871 (Collembola, Neanuridae) with a key to Iranian species of the genus’, Zootaxa, 5150 (3), 443–50.

Shayanmehr, Masoumeh and Elliyeh Yahyapour (2019), ‘The Collembola of North Forests of Iran, List of Genera and Species’, Journal of Environmental Science and Engineering B, 8 (4),

Shayanmehr, Masoumeh, Elham Yoosefi Lafooraki, and Morteza Kahrarian (2020), ‘A new updated checklist of Iranian Collembola (Arthropoda: Hexapoda)’, Journal of Entomological society of Iran, 39 (4), 403–45.

Shayanmehr, Masoumeh, et al. (2024), ‘New records of springtails (Hexapoda, Collembola) for Iran from the Bula Hyrcanian forest’, Journal of Insect Biodiversity and Systematics, 10 (1), 31–42.

Shayanmehr, Masoumeh, et al. (2025), ‘Entomological Society of Iran Faunal Analysis’, J. Insect Biodivers. Syst., 11 1085–1085.

Skarzynski, D and A Smolis (2006), ‘Skoczogonki [Collembola] rezerwatu’Srubita’w Beskidzie Zywieckim’, Parki Narodowe i Rezerwaty Przyrody, 25 (2), 41–50.

Skubała, Piotr and Anna Marzec (2013), ‘Importance of different types of beech dead wood for soil microarthropod fauna’, Polish Journal of Ecology, 61 (3), 545–60.

Sławska, M, A Bruckner, and M Sławski (2017), ‘Edaphic Collembola assemblages of European temperate primeval forests gradually change along a forest-type gradient’, European Journal of Soil Biology,

Sławski, Marek and Małgorzata Sławska (2020), ‘Collembolan Assemblages Response to Wild Boars (Sus scrofa L.) Rooting in Pine Forest Soil’, Forests, 11 (11), 1123.

Smolis, Adrian and Dariusz Skarżyński (2003), ‘Springtails (Collembola) of the” Przełom Jasiołki” reserve in the Beskid Niski Mountains (Polish Carpathians)’,

Söderström, L (1989), ‘Regional distribution patterns of bryophyte species on spruce logs in northern Sweden’, Bryologist,

Hopkin, Steve P. (2006), ‘Collembola’, in Lal, Rattan (ed.), Encyclopedia of Soil Science (1937.

Susanti, WI, et al. (2021), ‘Conversion of rainforest into oil palm and rubber plantations affects the functional composition of litter and soil Collembola’, Ecol Evol, 11 (15), 10686–708.

Thakur, MP, et al. (2017), ‘Warming magnifies predation and reduces prey coexistence in a model litter arthropod system’, Proc Biol Sci, 284 (1851), 20162570.

The R Core Team (2025), ‘R: A Language and Environment for Statistical Computing’, http://www.R-project.org/,

Theenhaus, A, S Scheu, and M Schaefer (1999), ‘Contramensal interactions between two collembolan species: effects on population development and on soil processes’, Functional Ecology, 13 (2), 238–46.

Tohidifar, M, et al. (2016), Biodiversity of the Hyrcanian Forests: A synthesis report, (UNDP/GEF/FRWO Caspian Hyrcanian Forest Project, Iran:) 41.

Tuomisto, Hanna (2010a), ‘A diversity of beta diversities: straightening up a concept gone awry. Part 2. Quantifying beta diversity and related phenomena’, Ecography, 33 (1), 23–45.

Tuomisto, Hanna (2010b), ‘A diversity of beta diversities: straightening up a concept gone awry. Part 1. Defining beta diversity as a function of alpha and gamma diversity’, Ecography, 33 (1), 2–22.

Tuomisto, Hanna (2011), ‘Commentary: do we have a consistent terminology for species diversity? Yes, if we choose to use it’, Oecologia, 167 (4), 903–11.

Vakili, Mehdi, et al. (2021), ‘Resistance and Resilience of Hyrcanian Mixed Forests Under Natural and Anthropogenic Disturbances’, Frontiers in Forests and Global Change, 4

Valle, Barbara, et al. (2023), ‘Finding the optimal strategy for quantitative sampling of springtails community (Hexapoda: Collembola) in glacial lithosols’, Pedobiologia, 101 150914.

Wickham, Hadley (2009), ggplot2: Elegant Graphics for Data Analysis, (Use R, New York: Springer-Verlag).

Widenfalk, Lina A, et al. (2016), ‘Small-scale Collembola community composition in a pine forest soil– Overdispersion in functional traits indicates the importance of species interactions’, Soil Biology and Biochemistry, 103 52–62.

Xie, Z, et al. (2022), ‘Drivers of Collembola assemblages along an altitudinal gradient in northeast China’, Ecol Evol, 12 (2), e8559.

Xie, Zhijing, et al. (2025), ‘Taxonomic and functional β-diversity of Collembola across elevational and seasonal gradients on a temperate mountain’, Geoderma, 458 117322.

Yahyapour, E, R Vafaei-Shoushtari, and M Shayanmehr (2019), ‘Study on the Collembola fauna of Mazandaran (Iran) and new records from the forests of this province’, Journal of Entomological Research, 10 (4), 103–13.

Yahyapour, E, et al. (2020), ‘A review of the Iranian species of the family Onychiuridae (Collembola, Poduromorpha), with description of five new species from Hyrcanian Forests in Iran’, Zootaxa, 4861 (1), zootaxa.4861.1.1.

Yahyapour, Eliye, et al. (2021), ‘New records of springtails (Hexapoda: Collembola) for Iran from Mazandaran forests’, Journal of Insect Biodiversity and Systematics, 7 (3), 263–76.

Yoosefi Lafooraki, Elham, et al. (2020), ‘Isotomidae (Collembola) from northern Iran with description of a new species of Isotomodes Linnaniemi’, Journal of Asia-Pacific Biodiversity, 13 (4), 545–53.

